# Stability Analysis of Delayed Age-structured Resource-Consumer Model of Population Dynamics with Saturated Intake Rate

**DOI:** 10.1101/2020.01.10.901629

**Authors:** Vitalii V. Akimenko

## Abstract

The nonlinear n-resource-consumer autonomous system with age-structured consumer population is studied. While the model of consumer population dynamics is described by a delayed transport equation, the dynamics of resource patches are described by ordinary differential equations with saturated intake rate. The delay models the digestion period of generalist consumer and is included in the calorie intake rate which impacts on consumer’s fertility and mortality. Saturated intake rate models the inhibition effect from the behavioural change of the resource patches when they react on the consumer population growing or from the crowding effect of the consumer. The model is studied both analytically and numerically. Conditions for existence of trivial, semi-trivial and non-trivial equilibria and their local asymptotic stability are obtained. Numerical experiments confirm and illustrate these theoretical results.

## 1. Introduction

Competition between several food patches and a common generalist consumer was thoroughly studied in the ecological literature [2], [23] - [27], [30] - [32], [34], [43], [44]. Here, an increasing in biomass of one food patch causes increasing in generalist consumer population, thus a negative impact on other resources patch and vice versa. Thus, apparent competition is similar to exploitative competition [33] and can lead for reducing the number of coexisting resource patches. In traditional unstructured Lotka-Volterra ODE models of population dynamics, many details of life histories are neglected [16]. The more reasonable approach in population dynamics modelling is based on the physiologically structured models [8], [11], [13], [14], [16] - [21], [28], [29], [35], [36], [41], [42]. In this article the apparent competition model of unstructured resource patches with age-structured consumer population is studied. This approach allows us to relate foraging to the life history and demographical characteristics of consumer population (fertility and mortality).

The dynamic interaction between resources and consumers in predator-prey models is described by the consumer’s functional response. Beddington [10] and DeAngelis [15] introduced and analysed the functional response with saturation which is often used now in applied models providing the more realistic description of predator-prey interaction [12], [22], [37], [40]. The functional response of such type, “saturated incidence rate”, was first introduced in SIR epidemic model after studying the cholera epidemic spread in Bari [12] and was used later in the various epidemic models [22], [40]. The feature of saturated incidence rate is that it tends to saturation when the population of predator (or infectives in epidemic models, parasite in parasite-host model, consumers in resource-consumer models, etc.) gets large and, as a consequence, it prevents the unboundedness of the contact rate between prey and predator. Since this functional response considers the behavioral change of prey (or hosts, susceptibles, resources in patches, etc.) as a reaction on the predator population growing or “crowding effect” of predator, the resource-consumer models with intake rates of such form are more reasonable in comparison with traditional Lotka-Volterra models. The resource consumption in biological and ecological models is often characterized also by the calorie intake rate which depends linearly from the amount of food resource taken by one consumer per unit of time from all patches. This function depends from the handling time, i.e. the time a consumer needs to handle and digest a unit of resource. This time period is included in model as a time delay parameter. Thus, the resulting model studied in this article consists of several unstructured resource patches and a single age-structured consumer population that forages in these patches including the saturated intake rate, calorie intake rate and the digestion period of a generalist consumer as a time delay parameter. The model is formulated in Section 2.

The conditions of existence of the trivial, semi-trivial and non-trivial equilibria of autonomous system are studied in Sections 3. The local asymptotic stability of all equilibria is considered in Section 4. Stability analysis is based on the traditional perturbation theory and linearization of autonomous system and includes the study of impact of the time delay parameter on the asymptotic stability of equilibria [3], [8], [19], [36], [38], [39].

The recurrent algorithms obtained in the earlier works [4] - [6] are used in the Section 5 for numerical analysis of dynamical regimes of autonomous system that were considered in the previous sections. In the first and second groups of experiments the local asymptotic stability of the trivial and semi-trivial (i.e., only resources exist at positive densities) equilibria for three resource patches with one generalist consumer is studied. Depending on the reproduction number of consumers, trajectories of system are unstable, oscillate in the vicinities of the trivial and semi-trivial equilibria or converge asymptotically to the semi-trivial equilibrium. The further increasing of consumer’s basic reproduction number or time delay parameter leads to the consumer population outbreaks in form of pulse sequence, which are classified in the quantitative population ecology as the populations with cyclical eruption dynamics [1], [7]. The results of simulations illustrated the properties of the outbreak solutions are presented in Section 5.2.

The next group of experiments focuses on the study of asymptotic behaviour of solutions in the vicinity of the non-trivial equilibrium with one non-depleted and two depleted resource patches (1^st^ - 3^th^ experiments), three non-depleted resource patches (4^th^ experiment) and one non-trivial consumer population. The results of simulations confirm and illustrate the statements of theorems and exhibit the different dynamical regimes of system with unstable and asymptotically stable trajectories for the selected parameters of the model. Several concluding remarks are given in the last Section 6.

## 2 Model

In this article we study an apparent competition food web module that consists of *n* resource patches and consumers that move freely between these patches. Resource density of *i* -th patch is denoted as *y*_*i*_(*t*), *i* = 1,…, *n*. The resource dynamics in each patch is described by the logistic model with constant growth rate *r*_*i*_ > 0, and environmental carrying capacity *K*_*i*_ > 0. The age-specific density of consumer population at age *a* and time *t* is denoted by *w*(*a,t*), the quantity of consumers in population is 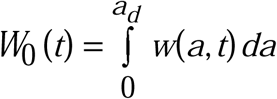 (where *a*_*d*_ > 0 is the maximum consumer’s life-span) and a weighted quantity of consumers at the fixed time *t* is 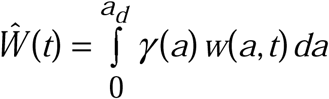 (where *γ*(*a*) is an age-specific consumer’s preferences in food resource, *γ*(*a*)∈(0,1], *γ*(*a*)∈*L*_2_ ([0, *a*_*d*_])). The interaction strength between resources and consumers is a product of the saturated intake rate *y*_*i*_(*t*)*g*_*i*_(*Ŵ*(*t*)), where *g*_*i*_(*Ŵ*(*t*)) evolves to a saturation level when *Ŵ*(*t*) gets large, i.e. *g*_*i*_(*Ŵ*(*t*)) = *β*_*i*_*Ŵ*(*t*)(1 + *α*_*i*_*Ŵ*(*t*))^−1^. Functions *g*_*i*_(*Ŵ*(*t*)) have a form of the Beddington – DeAngelis type of functional responses [10], [15], [37] under assumption that handling time of predator is effectively zero (Eq. (12) in [10]). Constant *β*_*i*_ > 0 is a search rate of resource *i* = 1,…, *n*. Saturation coefficient *α*_*i*_ ≥ 0 is proportional to the rate of encounter between consumers, related both to their speed of movement and the range at which they sense each other and the time wasted by consumer per one encounter [10]. On the other hand, this coefficient can consider also the behavioral change of the food resource when consumer population grows (like in epidemic models for pair susceptibles-infectives [12], [22], [40]). The greater the coefficient *α*_*i*_, the greater the activity of the food resource in *i* –th patch and vice versa. When *α*_*i*_ = 0 the saturated intake rate is a bilinear form of Lotka-Voltera functional response which considers the inactive food resource without behavioral reaction on the consumer population changes. For our convenience we introduce the food resource classification: the higher activity resource with 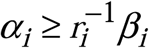, the lower activity resource with 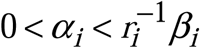 and non-active resource with *α*_*i*_ = 0. These assumptions lead to the following resource population dynamics

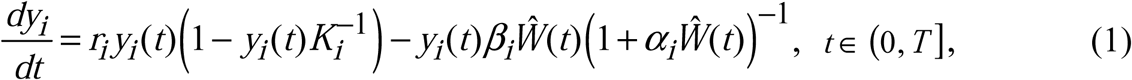

Consumer population dynamics *w*(*a,t*) are governed by the delayed McKendrick-Von Foerster’s age-structured model [18], [20], [21]:

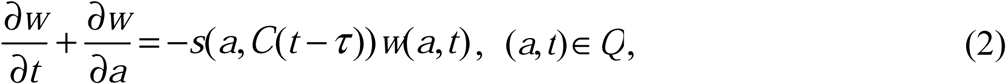

where 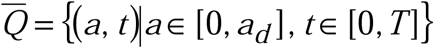. Eqs. (1), (2) are completed by the following initial and boundary conditions:

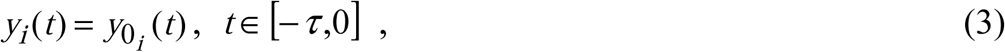

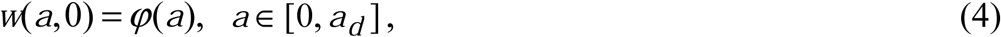

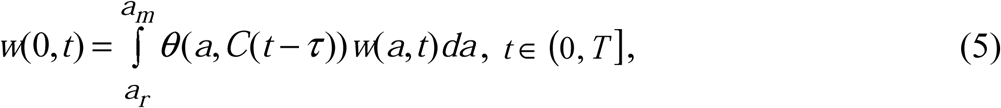

where *a*_*r*_ > 0 is an age of maturation, *a*_*m*_ > 0 is a maximum age of reproduction and *φ*(*a*) is an consumer’s initial density. Functions *s*(*a,C*(*t* − *τ*)) and *θ*(*a,C*(*t* − *τ*)) are age- and calorie intake rate dependent consumer’s death and fertility rates, respectively. Consumption of food resources by one consumer per unit of time is measured by calorie intake rate *C*(*t*). This function is used in Eqs. (2), (5) with the time delay parameter *τ* > 0 which is a handling time, i.e. the time a consumer needs to handle and digest a unit of resource. We assume that the calorie intake rate is a linear function of the amount of food resource taken by one consumer per unit of time from all patches and is defined through the resource intake rate:

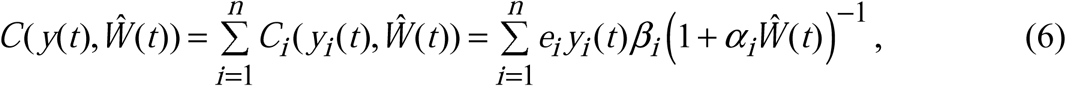

where *C*_*i*_ ≥ 0 is a calorie intake rate for *i* -th resource patch, *e*_*i*_ > 0 is an efficiency with which the consumed resource *i* is transformed to energy. We impose the following natural restrictions on the death and fertility rates:

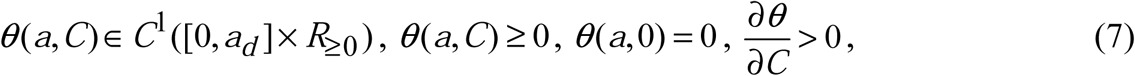

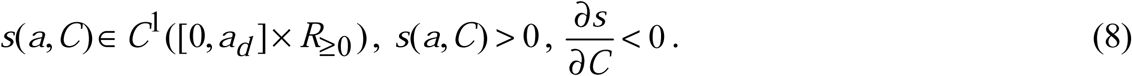

Equations (7), (8) means that decreasing of calorie intake rate corresponds to the critical foraging or starvation, and increasing of it corresponds to the sufficient foraging and satiation with increasing resource intake rate. Increasing of calorie intake rate provides also maximum comfortable conditions for reproduction of consumers that corresponds to increasing of birth rate, and decreasing of calorie intake rate provides the most poor and unfavourable conditions for reproduction of consumers, decreasing of birth rate.

The basic reproduction number of age-structured model of consumer population dynamics [8], [16], [28], [35], [41] is a calorie intake rate depending function:

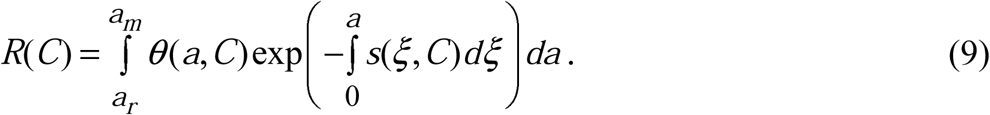

## 3 Existence of stationary equilibria of autonomous system (1) – (5)

We consider the equilibria 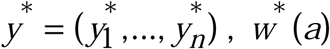, *w*^*^(*a*) of autonomous system (1) – (5). It is easy to verify that trivial *y*^*^ = (0,…,0), *w*^*^(*a*) ≡ 0 and semi-trivial *y*^*^ = (*K*_1_,…, *K*_*n*_), *K* = (*K*_1_,…, *K*_*n*_), *w*^*^(*a*) ≡ 0, equilibria of autonomous system (1) – (5) with coefficients (7), (8) always exist. In this section we study the conditions for existence of a nontrivial equilibrium where resource densities in the group of patches with indexes *i*∈ *I*_+_ are positive and bounded 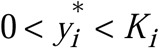, while the remaining patches with indexes *i*∈ *I*_0_ are depleted 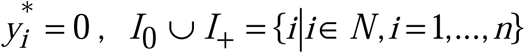. The equilibrium consumer density is nonnegative *w*^*^(*a*) ≥ 0, equilibrium consumer quantity and equilibrium weighted consumer quantity are positive 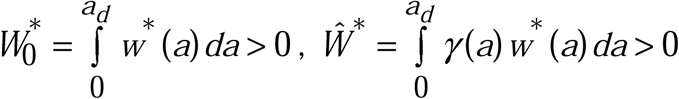. Equilibrium 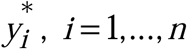, satisfies equation:

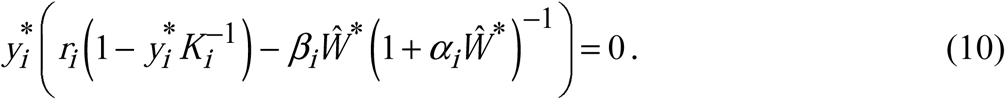

Equation (10) has at most two nonnegative solutions:

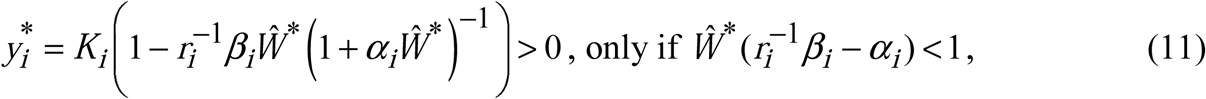

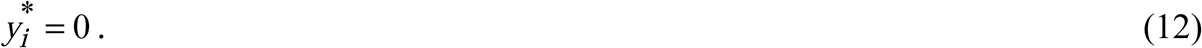

Hence, the nontrivial equilibrium of food web contains the nonempty set of nondepleted patches which necessarily satisfy condition 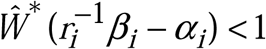 with indexes 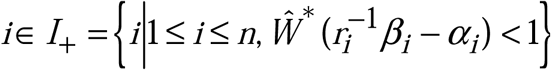 and the set (empty or not) of depleted patches which can satisfy or not the condition 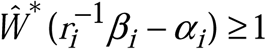 with indexes *i*∈ *I*_0_ : *I*_0_ ⋃ *I*_+_ ={*i*|*i*∈ *N, i*= 1,…, *n*}, *I*_0_ ⋂ *I*_+_ = Ø.

From Eq. (11) we obtain the positive equilibria *Ŵ* ^*^ > 0 :

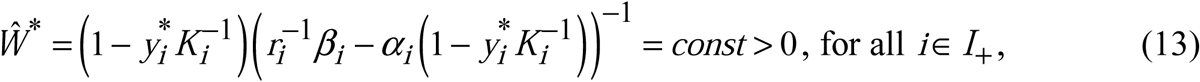

or

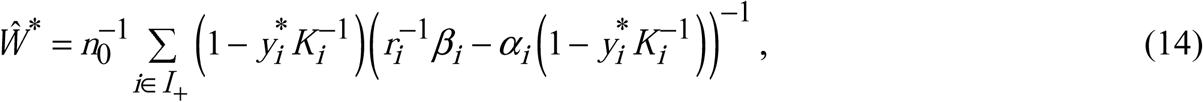

where *n*_0_ is a number of patches of set *I*_+_ ≠ Ø, 1 ≤ *n*_0_ ≤ *n*. Substituting Eqs. (11) and (13) in Eq. (6) we obtain the equilibrium calorie intake rate:

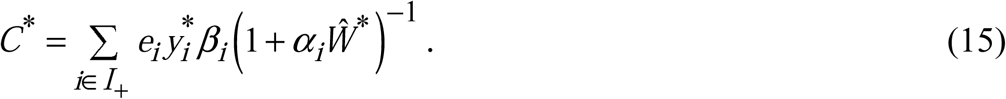

The second equation of equilibrium is obtained from Eqs. (2), (5):

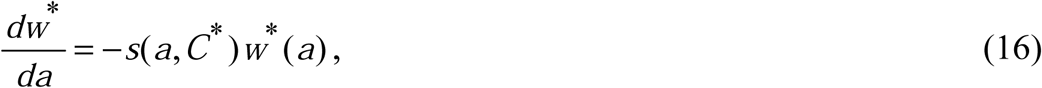

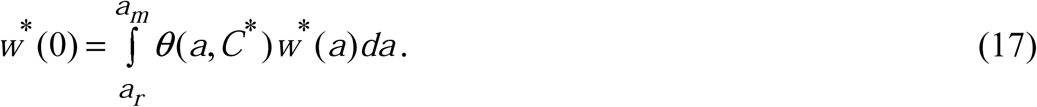

Solution of (16), (17) satisfies the following integral equation:

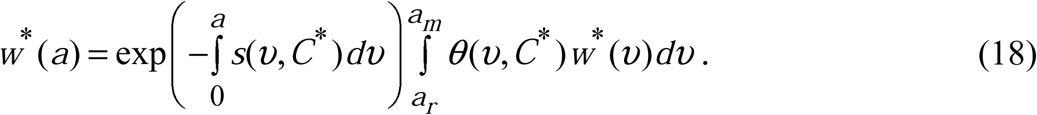

Integrating Eq. (18) with respect to *a* from 0 to *a*_*d*_ we obtain the expression with 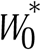:

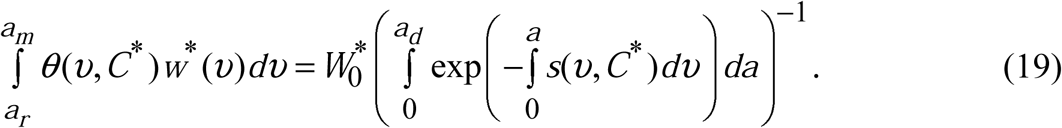

Multiplying both sides of Eq. (18) on *γ*(*a*), integrating them with respect to *a* from 0 to *a*_*d*_ and substituting in obtained equation the left side of Eq. (19) yields:

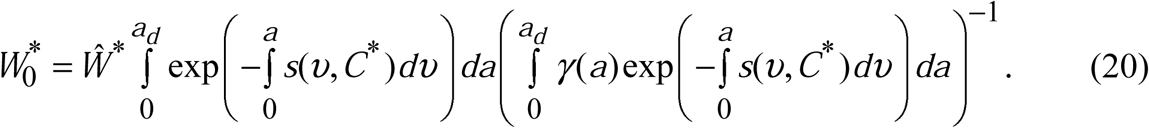

By analogy with Theorem 1 from [29] formulated for harvesting problem, we obtain

### Theorem 1.

The system (1) – (5) possess a nontrivial equilibrium 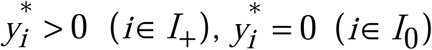, and 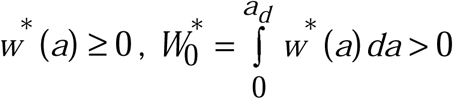, if and only if there exists the positive solution *Ŵ* ^*^ > 0 of equation *R*(*C*(*y*^*^(*Ŵ* ^*^),*Ŵ* ^*^)) = 1 with restrictions 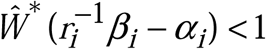, *i*∈ *I*_+_. The basic reproduction number *R*(*C*^*^), equilibrium 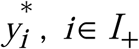, equilibrium calorie intake rate *C*^*^ and 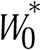 are given by Eqs. (9), (11), (15), and (20), respectively. The equilibrium distribution of consumer’s density is:

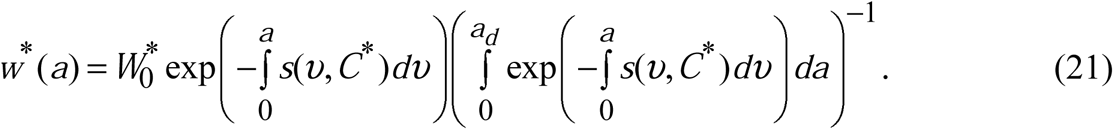

**Proof**. Multiplying both sides of Eq. (18) on *θ*(*a,C*^*^), integrating them with respect to *a* from *a*_*r*_ to *a*_*m*_ after a little algebra we arrive to the equation *R*(*C*^*^(*y*^*^(*Ŵ* ^*^),*Ŵ* ^*^)) =1 (see Eq. (9)). If this equation has solution *Ŵ* ^*^ > 0 satisfied 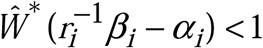 (*i*∈ *I*_+_, see Eq. (11)), 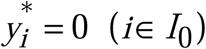, we can obtain 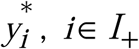, (Eq.(11)), *C*^*^(Eq.(15)) and 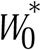 (Eq. (20)). Conversely, if equation *R*(*C*(*y*^*^(*Ŵ* ^*^),*Ŵ* ^*^)) = 1 does not have solution *Ŵ* ^*^ > 0 satisfied 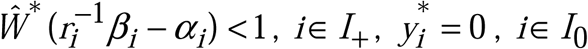, the stationary solution of problem (11), (12), (18), (20) does not exist.

Substituting the left-hand side of Eq. (19) in Eq. (18) we obtain the equilibrium distribution of consumer’s density (21). Theorem 1 is proved.

### Corollary 1.

If the sets of indexes of the lower and higher activity resourcessome patches have higher activity resources with 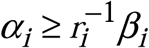, condition 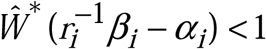 holds for them and such patches always have positive equilibria 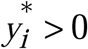 defined by Eq. (11) (i.e. non-depleted patches).

Theorem 1 imposes the restriction on the basic reproduction number of consumer population *R*(*C*(*y*^*^,*Ŵ* ^*^)) at the equilibrium *y*^*^, *Ŵ* ^*^ taking into account the impact of foraging on the consumer fertility and mortality. The condition of existence of nontrivial balance between food resource growing and consumer demographical processes (nontrivial equilibrium) is given in the form of transcendental integral equation. Implementation of such condition in biological applications is difficult from the technical point of view. In the next theorem we provide the sufficient conditions on coefficients of the system (1) - (5) that guarantee existence of the nontrivial equilibrium.

### Theorem 2.

Let the sets of indexes of the lower and higher activity resources of non-depleted patches are 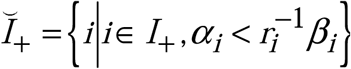 and 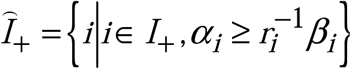, respectively, 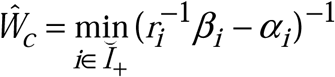,

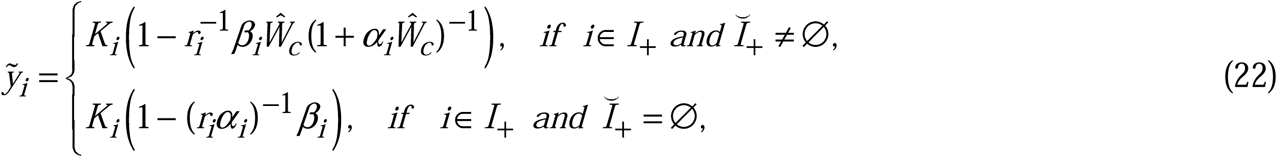

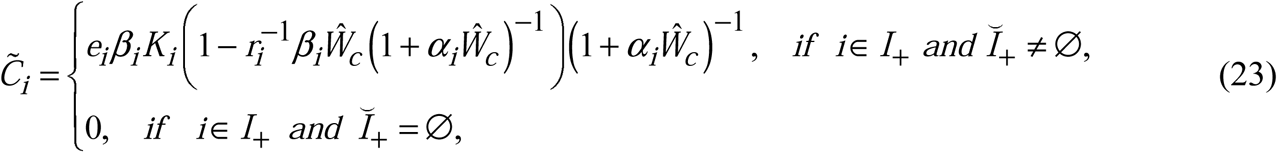

and coefficients of system (1) - (5) satisfy Eqs. (7), (8). Then, for existence of at least one non-trivial solution of stationary problem (11), (12), (18) – equilibrium 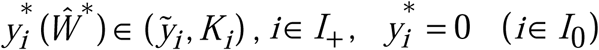 and 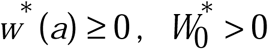 it is sufficient that 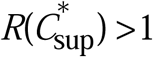 and 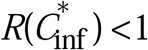 where the infimum and supremum of equilibrium calorie intake rate are:

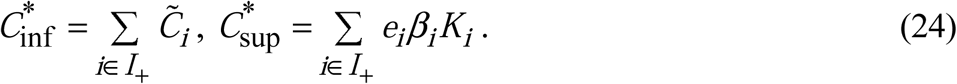

**Proof**. From Eq. (11) we obtain the partial derivative of 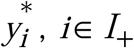, at *Ŵ* ^*^ > 0 :

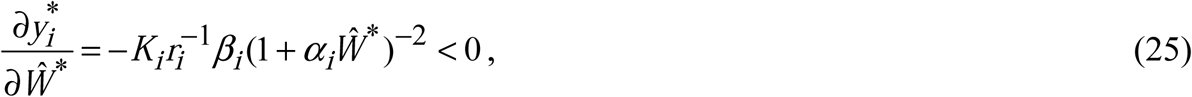

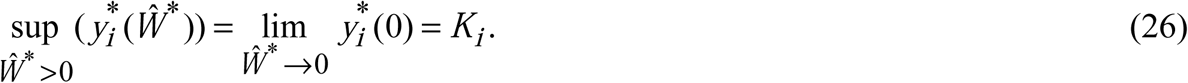

For the higher activity non-depleted resources 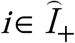 equilibrium 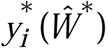 Eq. (11)) is positive at *Ŵ* ^*^ > 0 and its infimum is:

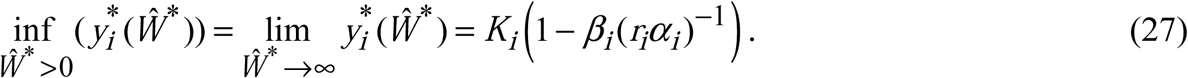

For the lower activity non-depleted resources 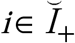 equilibrium 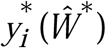 is positive only at 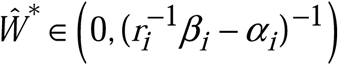 and its infimum is:

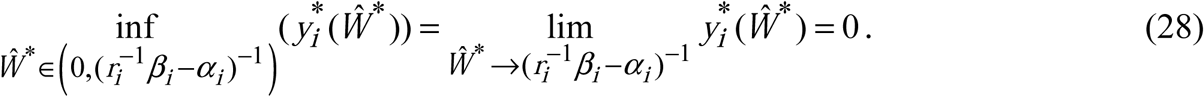

From Eqs. (26) - (28) we obtain the supremum of *Ŵ*^*^ for which the restriction 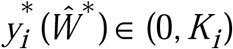 holds for all 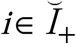:

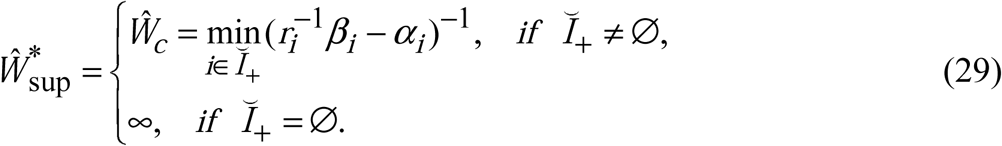

From Eq. (29) we obtain the infimum of 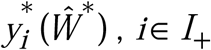, for all non-depleted patches:

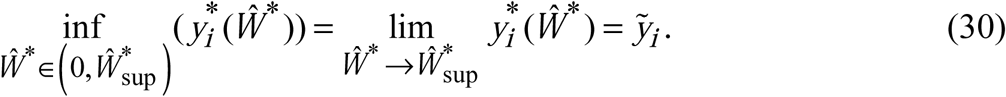

Hence, from Eqs. (26), (30) it follows that 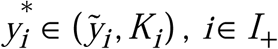. Substituting Eq. (11) in Eq. (15) we obtain the equilibrium calorie intake rate of the form:

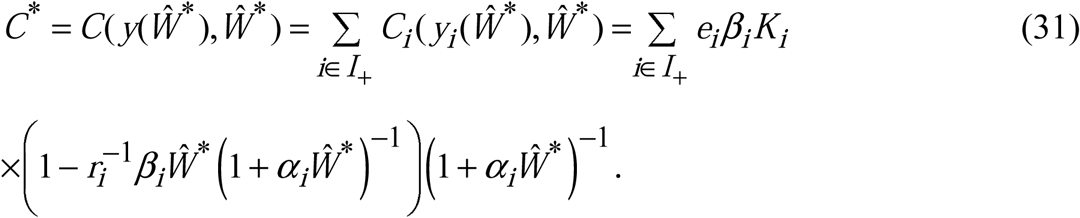

The partial derivative of *C*(*y*(*Ŵ* ^*^),*Ŵ* ^*^) at 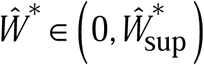 is:

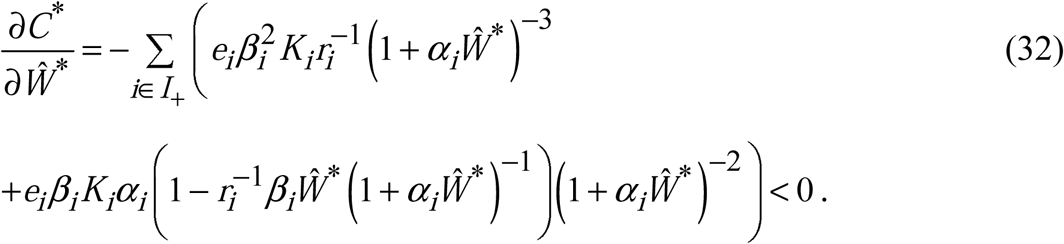

Using Eqs. (29), (31), (32) and restriction *C*_*i*_ (*y*_*i*_ (*Ŵ* ^*^),*Ŵ* ^*^) ≥ 0 we can estimate the infimum and supremum of *C*^*^:

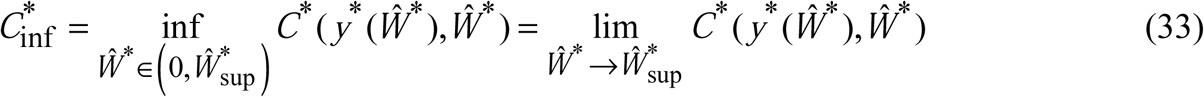

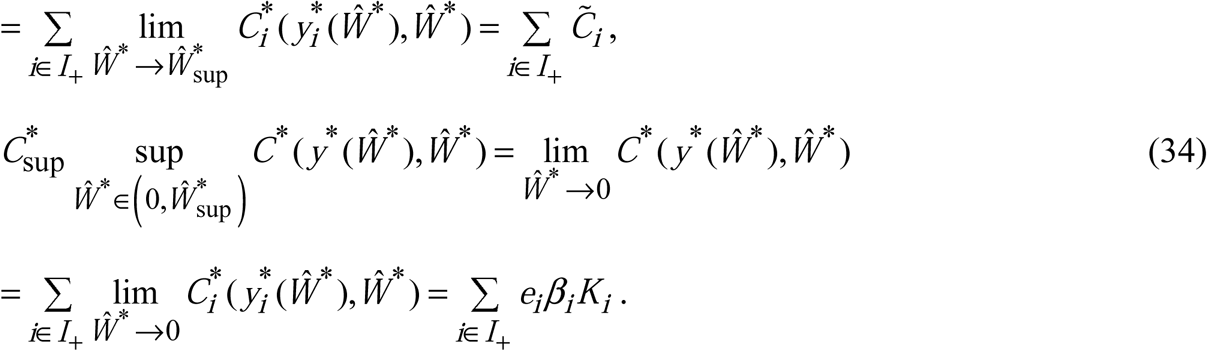

where 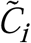 are given by Eq. (23). Taking into account the properties of death and fertility rates (7), (8), we obtain the derivative of basic reproduction number at 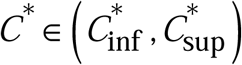:

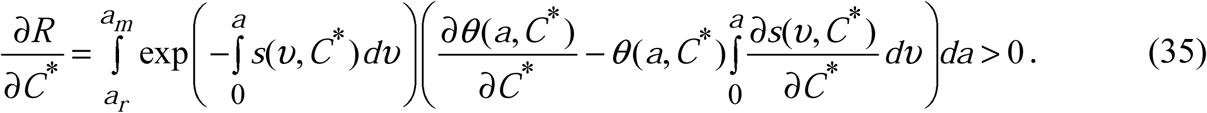

Hence, using Eqs. (33), (34), we can estimate the supremum and infimum of *R*(*C*^*^):

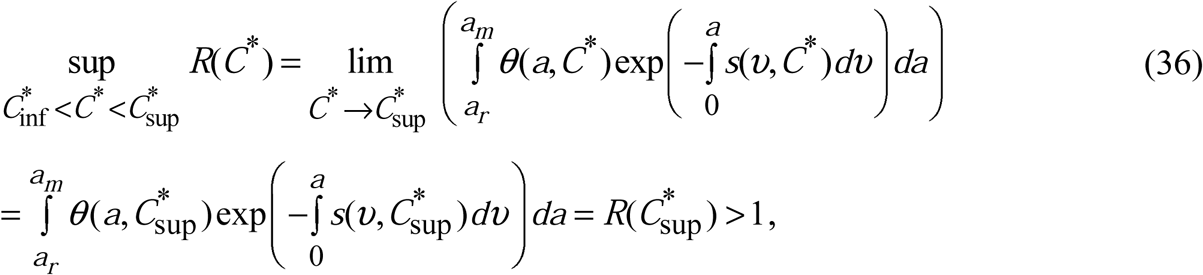

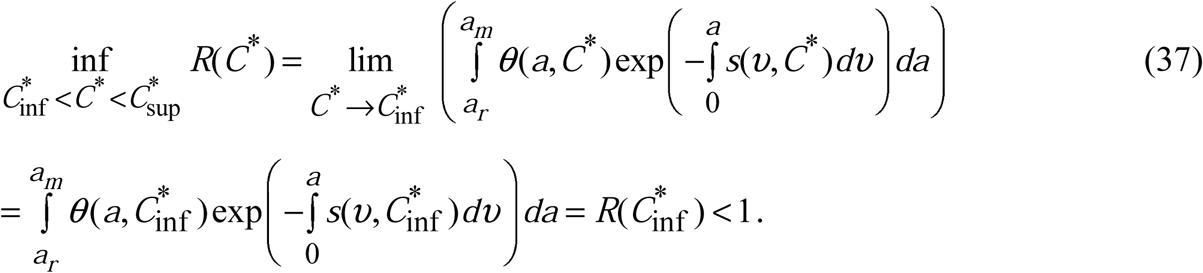

Since *R*(*C*(*y*^*^(*Ŵ* ^*^),*Ŵ* ^*^)) and *C*(*y*^*^(*Ŵ* ^*^),*Ŵ* ^*^) are continuous functions of 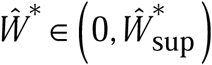, from Eqs. (36), (37) it follows that equation *R*(*C*(*y*^*^(*Ŵ* ^*^),*Ŵ* ^*^)) = 1 always has at least one positive solution *Ŵ* ^*^ > 0 for which 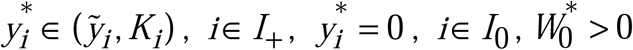, and *w*^*^(*a*) ≥ 0. Theorem 2 is proved.

### Corollary 1.

If all non-depleted patches have only lower activity resources with 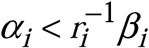, when 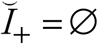, from Eq.(23) it follows that the infimum of equilibrium intake rate 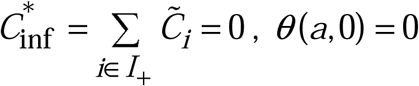 (Eq.(7)) and, consequently, *R*(0) = 0. In this case condition 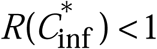 is always satisfied and can be omitted in Theorem 2.

## 4 Local asymptotic stability of equilibria of autonomous system (1) – (5)

### The asymptotic stability of trivial and semi-trivial equilibria

Linearizing Eqs. (1) at the trivial equilibrium 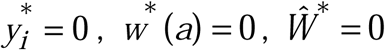, we arrive to the equation for perturbations *ζ* _*i*_(*t*) :

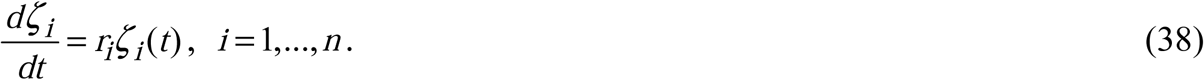

Since *r*_*i*_ > 0, the trivial equilibrium is always unstable. Linearizing system (1) – (5) in the vicinity of the semi-trivial equilibrium 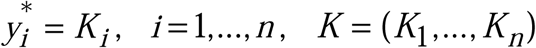, *w*^*^(*a*) = 0, we arrive to system for the perturbations *ζ* _*i*_(*t*) and *ξ*(*a,t*) :

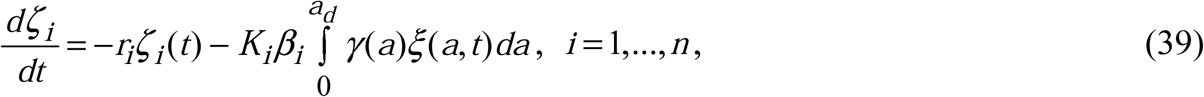

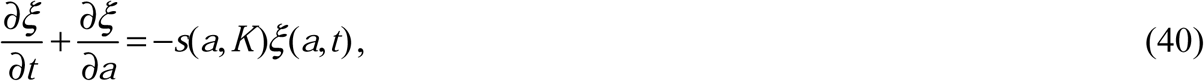

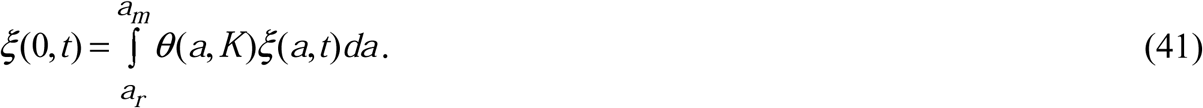

We consider solutions of system (39) – (41) in the form 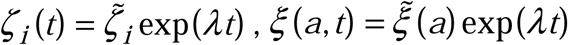:

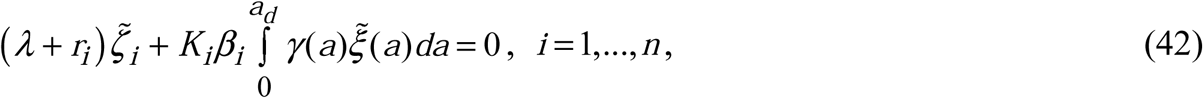

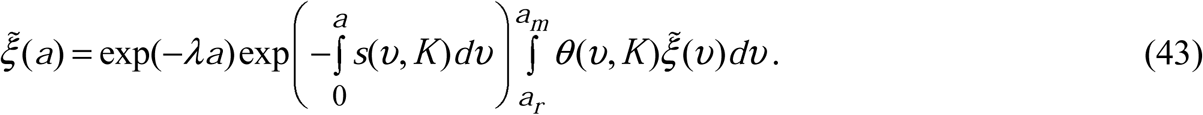

Multiplying Eq. (43) on *θ*(*a, K*) and integrating it with respect to *a* from *a*_*r*_ to *a*_*d*_ we arrive to the integral characteristic equation for unknown *λ*:

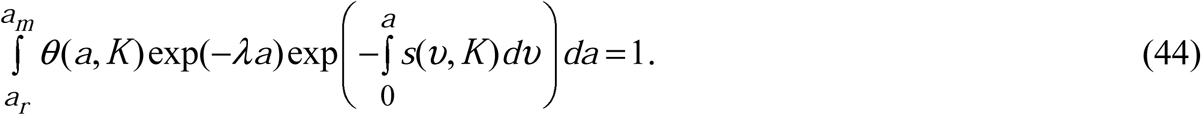

From Eq. (44) it follows that if the consumer’s reproduction number *R*(*K*) < 1, characteristic Eq. (44) has only negative root *λ*^*^ < 0. In this case perturbations *ξ*(*a,t*) → 0, and, consequently, *ζ* _*i*_(*t*) → 0 (see Eq. (39)). The semi-trivial equilibrium 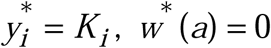 is locally asymptotically stable.

If *R*(*K*) = 1, Eq. (44) has only trivial root *λ*^*^ = 0 and the semi-trivial equilibrium is neutrally stable [38] (that is all solutions from the vicinity of semi-trivial equilibrium are unstable, but they do not have a form of asymptotic exponential growth). Finally, if *R*(*K*) > 1, the root of Eq. (44) is always positive *λ*^*^ > 0 and the semi-trivial equilibrium is unstable. Since Eqs. (42) - (44) do not depend from delayed parameter, the results obtained in this section are valid for all *τ* > 0. Now we can formulate the following theorem.

#### Theorem 3.

i. The trivial equilibrium 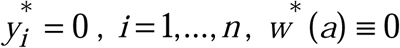 of system (1) – (5) is unstable for all *τ* > 0.
ii. The semi-trivial equilibrium of food resource 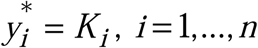, with *w*^*^(*a*) ≡ 0 is unconditionally (i.e. for all *τ* > 0) locally asymptotically stable if the consumer’s basic reproduction number *R*(*K*) <1 whereas it is unstable for all *τ* > 0 if *R*(*K*) ≥1.

#### Remark 1.

If consumer population is fully extinct and cannot renew the reproduction Eq. (43) has only the trivial solution 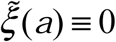. The roots of Eq. (42) are always negative *λ*^*^ = −*r*_*i*_ < 0, that is the perturbations *ζ*_*i*_(*t*) → 0 and the semi-trivial equilibrium is locally asymptotically stable. The examples of the nonlinear age-structured models of population dynamics with full extinction in the form of a single pulse population outbreak were obtained in works [6], [7].

### 4.2 Local asymptotic stability of nontrivial equilibrium

Linearizing system (1) – (5) at the nontrivial equilibrium 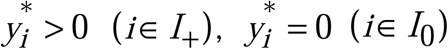, and 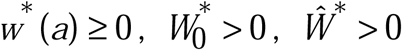 with perturbations 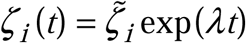 for 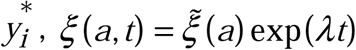 for *w*^*^ (*a*), and 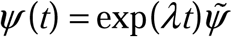 for *Ŵ* ^*^, where 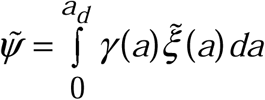, we arrive to the system:

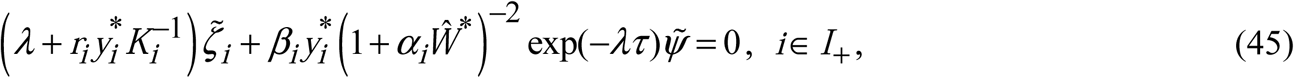

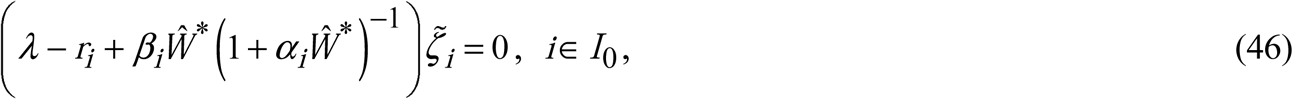

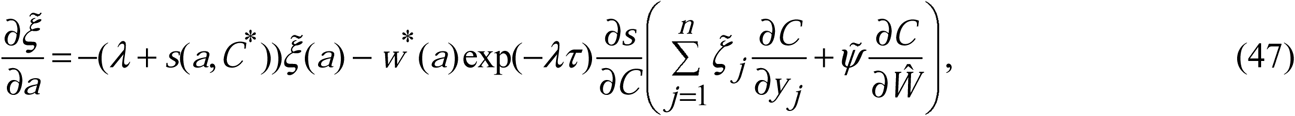

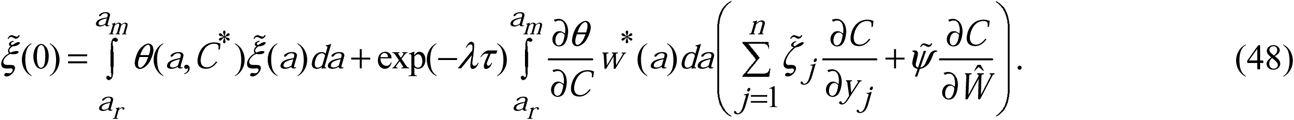

Integrating the linear ODE (47) with initial condition (48) we obtain the Fredholm integral equation of second type for 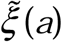:

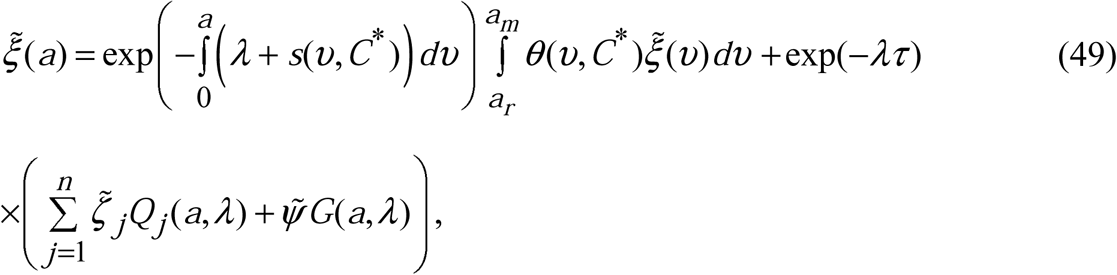

where

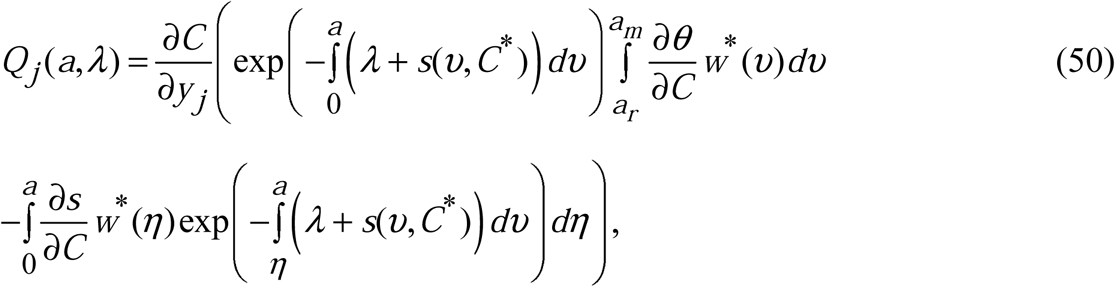

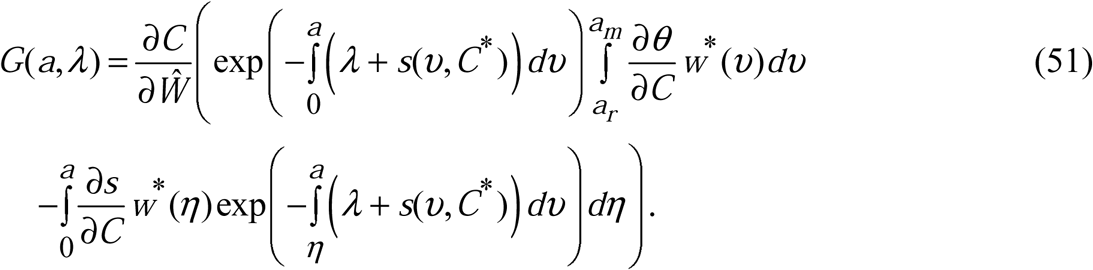

First, we analyse the existence of trivial root *λ* = 0 of linear system (45), (46), (49). Multiplying both sides of Eq. (49) on *θ*(*a,C*^*^), integrating them with respect to *a* from *a*_*r*_ to *a*_*d*_ and excluding *R*(*C*^*^) =1 from obtained expression yields:

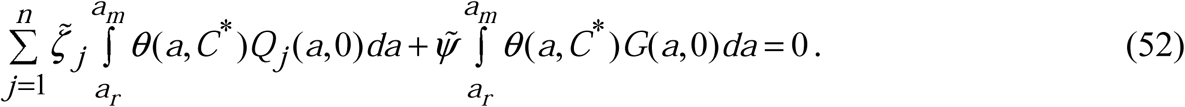

We arrive to the homogeneous linear system (45), (46), (52) of *n* + 1 order for the functions 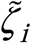 and 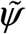. The determinant of this system is:

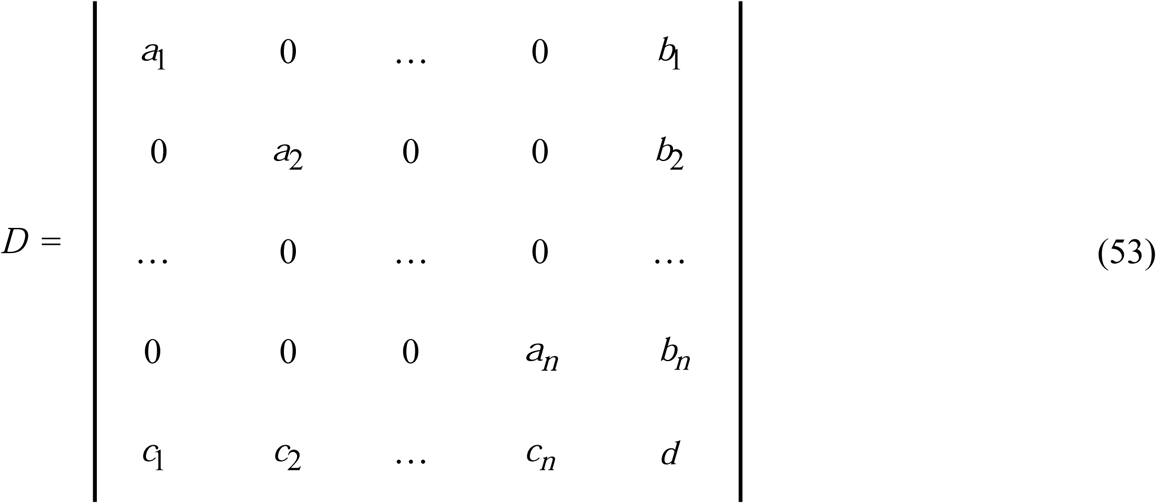

Here

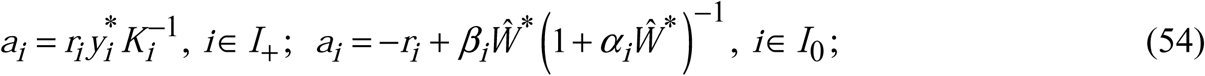

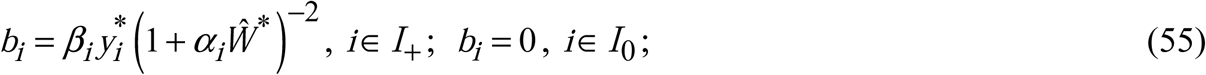

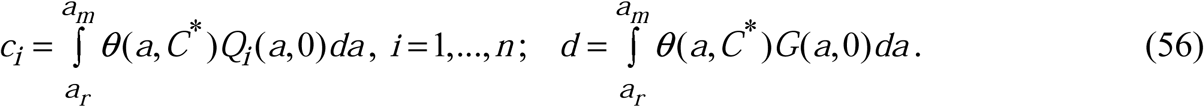

Equating to zero determinant *D* = 0 (53) we arrive to the characteristic equation:

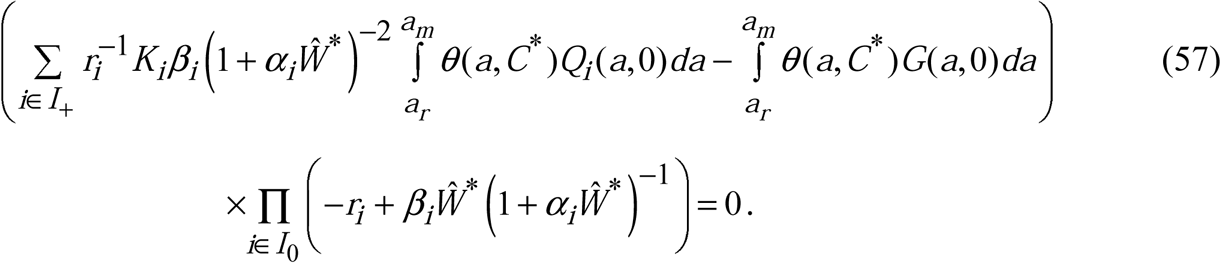

In Eq. (57) we assume that if equilibrium does not contain the depleted patches (*I*_0_ = Ø) the second multiplier of equation is omitted. If *I*_0_ ≡ Ø this multiplier equals zero only if there exists at least one depleted patch satisfied condition 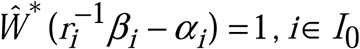. In this case is unstable. *λ* = 0 is a root of characteristic equation and the nontrivial equilibrium is unstable.

If 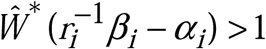 for all *i*∈ *I* _0_ the second multiplier in Eq. (57) is always negative and we have to analyse the first multiplier in brackets. The partial derivatives of the calorie intake rate (6) at the equilibrium *y*^*^, *Ŵ* ^*^ are:

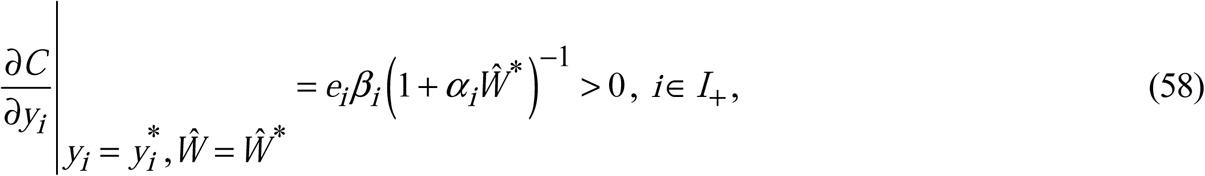

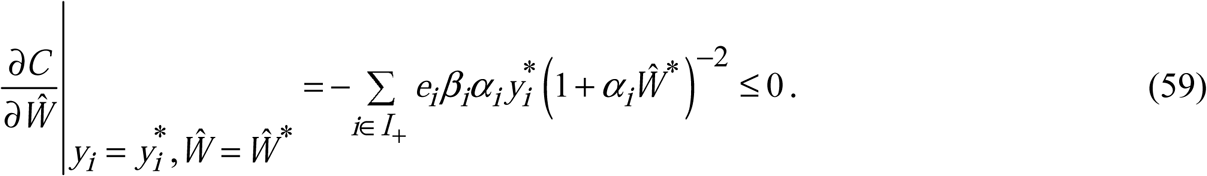

In Eq. (59) the partial derivative 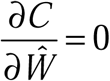 if all resources of non-depleted patches are non-active, i.e. *α*_*i*_ = 0 for all *i*∈ *I*_+_. From assumptions (7), (8) and Eqs. (50), (51), (58), (59) it follows that for *λ* = 0 the auxiliary functions have the properties:

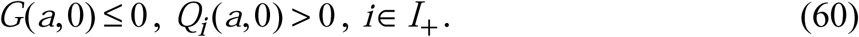

Then, the first multiplier (in brackets) in Eq. (57) is always positive, the left side of the equation is negative, and *λ* = 0 is not a root of characteristic equation (57). Thus, *λ* = 0 can be a root of Eq. (57) if and only if there exists at least one depleted patch with a lower activity resource for which 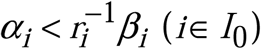 and 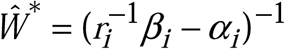.

Second, if *λ* ≠ 0 we can multiply both sides of integral equation (49) on *θ*(*a,C*^*^), integrate them with respect to *a* from *a*_*r*_ to *a*_*m*_ and substituting the obtained expression in Eq. (49) obtain the final solution of this equation:

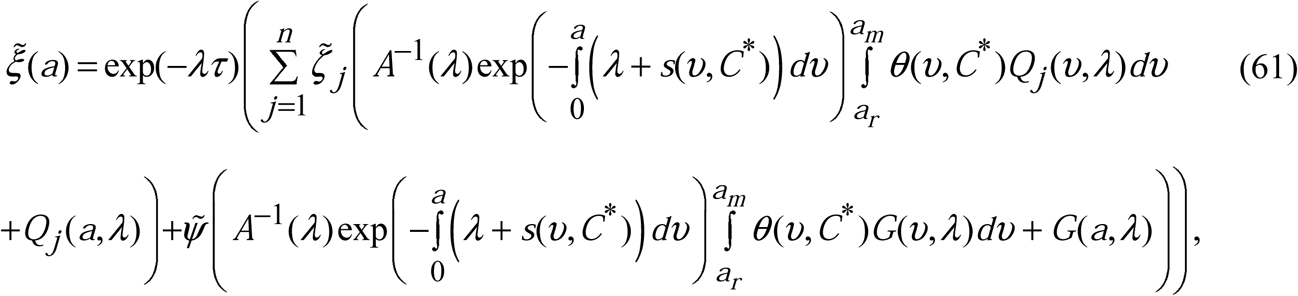

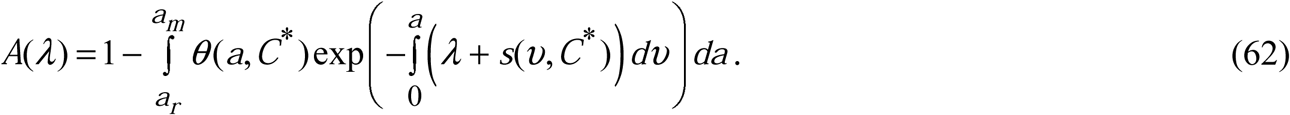

If *λ* ≠ 0 coefficient *A*(*λ*) ≠ 0 and solution (61) exists. For *λ* > 0 function *A*(*λ*) ∈(0,1). Multiplying both sides of Eq. (61) on *γ*(*a*), integrating them with respect to *a* from 0 to *a*_*d*_ yields:

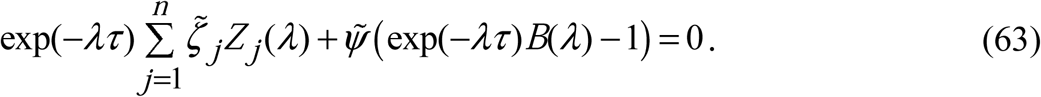

where the auxiliary functions *Z* _*j*_ (*λ*) and *B*(*λ*) are:

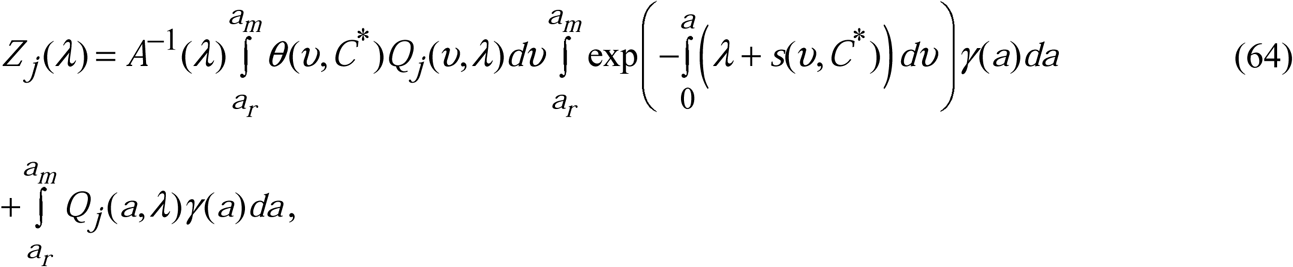

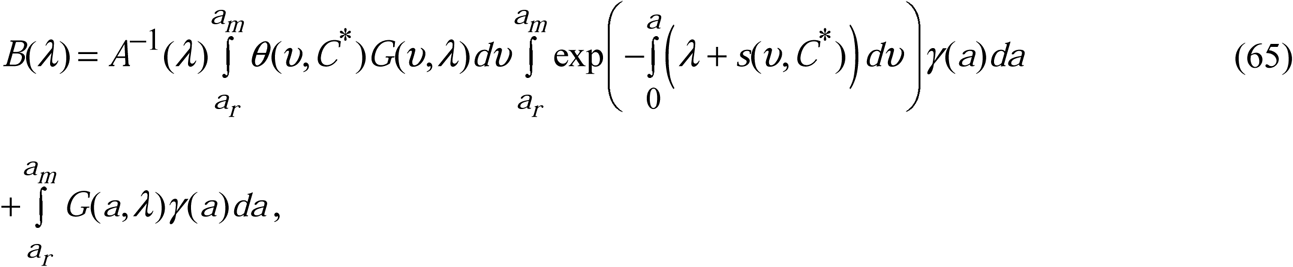

and the auxiliary functions *Q*_*j*_ (*a, λ*), *G*(*a, λ*), *A*(*λ*) are given by Eqs. (50), (51), (62), respectively. We arrive to the homogeneous linear system of *n* + 1 order (45), (46), (63) for the functions 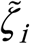 and 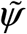. The determinant *D* of this system is given by (53), where

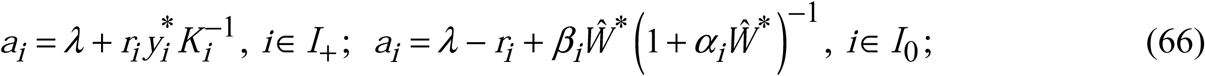

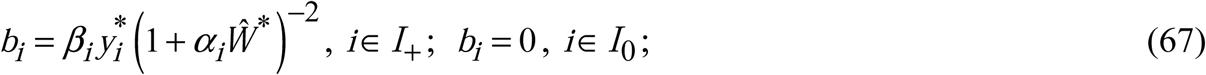

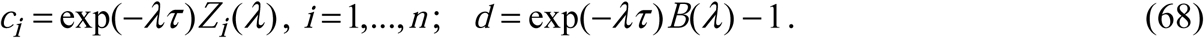

The characteristic equation of the system (45), (46), (63) is:

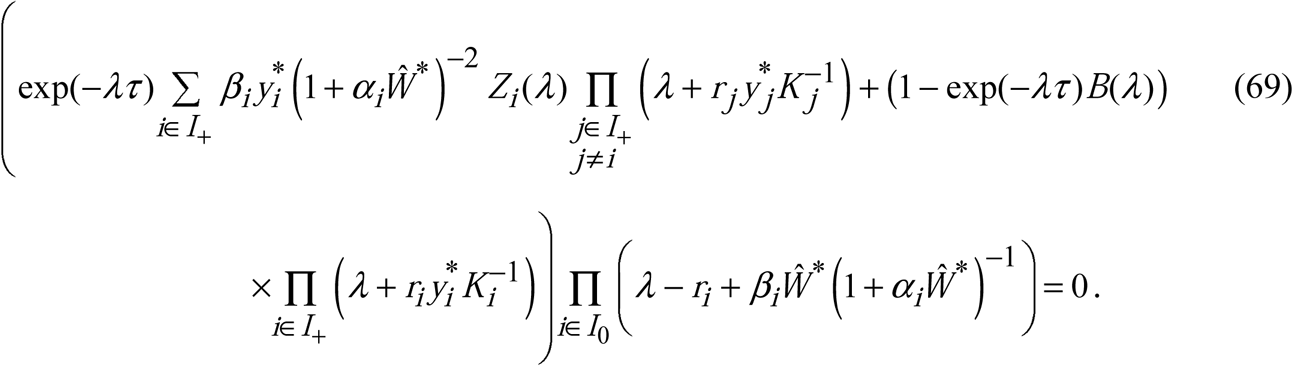

If *I*_0_ ≡ Ø, Eq. (69) has the real roots

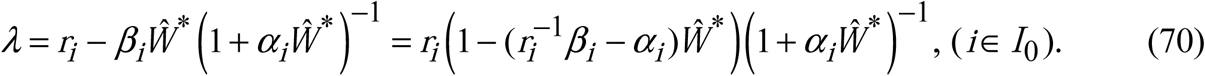

If at least one depleted patch has higher activity resource: 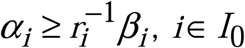, the corresponding root *λ* > 0 for any *Ŵ* ^*^ > 0 and the nontrivial equilibrium is unstable.

If all depleted patches have lower activity resources: 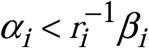 for all *i*∈ *I*_0_, and equilibrium 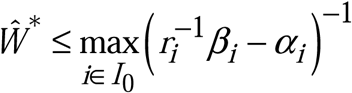, at least one of these roots *λ* ≥ 0 and the nontrivial equilibrium is unstable.

If 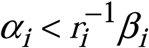 and 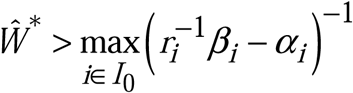, all roots in Eq.(70) are negative *λ* < 0, and for analysis of unstable equilibria we have to consider the others roots of Eq. (69) in the first brackets (non-depleted patches with *i*∈ *I*_+_). We have to include in this analysis also the case with only non-depleted patches, when *I*_0_ = Ø.

If Eq. (69) has a root 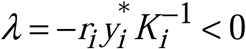 for some *i*∈ *I*_+_, the nontrivial equilibrium is asymptotically stable. Further we will assume that 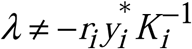 for all *i*∈ *I*_+_. Dividing the first brackets of Eq. (69) on 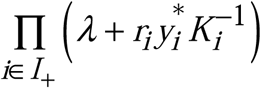 and equating it to zero we have:

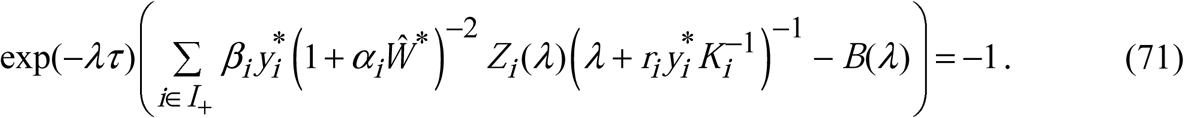

Since the partial derivatives of the calorie intake rate satisfy Eq. (58), (59) all auxiliary functions in Eq. (71) possess the properties:

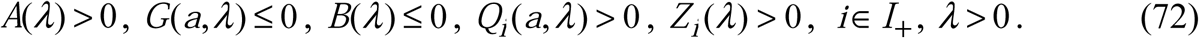

Hence, the left side of Eq. (71) is always positive for any time delay *τ* > 0 and *λ* > 0, and Eq. (71) does not have the real positive roots.

We have to analyse further the existence of complex roots of Eq. (69) with nonnegative real part when characteristic equation does not have real nonnegative roots, i.e. when *I*_0_ ≠∅ and 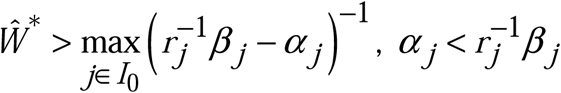, or when *I*_0_ = ∅. We seek the pure imaginary roots *λ* = *iω* (neutral stability state [38], where *ω* ≠ 0 is unknown real parameter) for which there exists the non-trivial solution of system (45), (46), (63). Substituting *λ* = *ω* in Eq. (46) and separating real and imaginary parts yields (*j* ∈ *I*_0_):

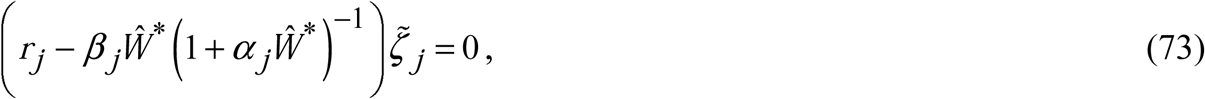

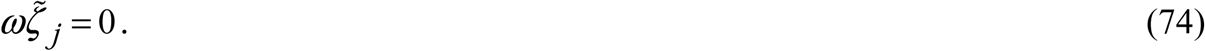

in Eq. (46)

Since 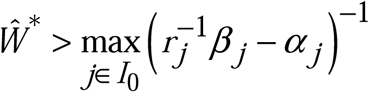 and 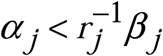, from Eq. (73) we obtain only the trivial solution for perturbations 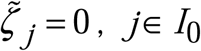 (the same result, when *I*_0_ =∅). Substituting *λ* = *iω* with 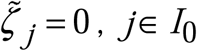, in Eq. (48) and separating real and imaginary parts yields:

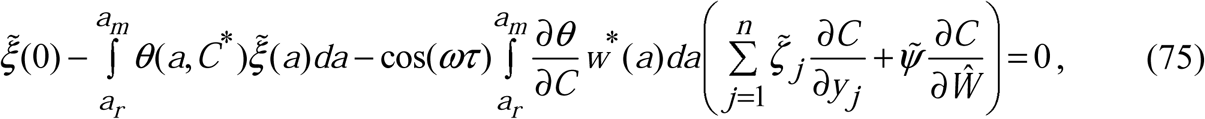

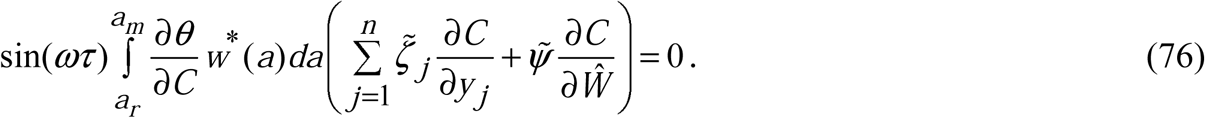

From Eqs. (7) and (76) it follows that nontrivial solution 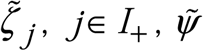 exists only if sin(*ωτ*) = 0. Indeed, if this is not true, we have

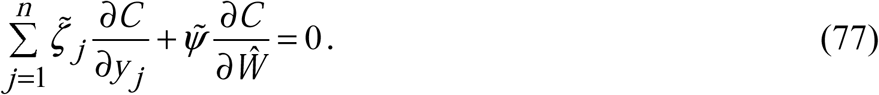

Substituting *λ* = *iω* in Eq. (47) and separating real and imaginary part yields:

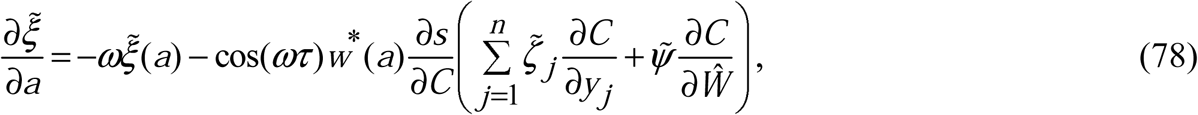

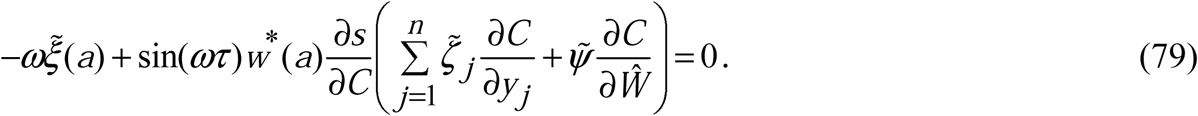

Multiplying Eq. (79) on *γ*(*a*) > 0 and integrating it with respect to *a* from 0 to *a*_*d*_, after a little algebra we arrive to the equation:

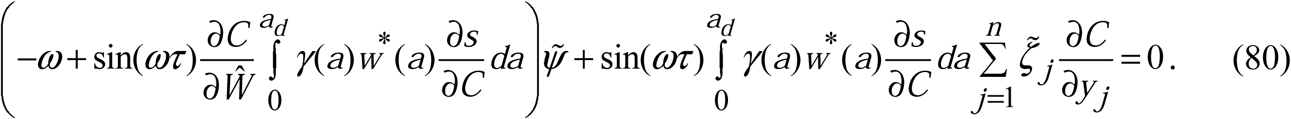

Substituting 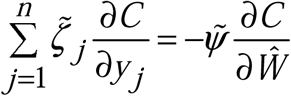from Eq. (77) in Eq. (80) we arrive to the equation with unique trivial solution:

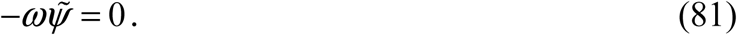

Hence, the nontrivial solution 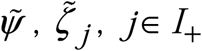 can exist only if Eq. (77) is not valid and sin(*ωτ*) = 0, cos(*ωτ*) = 1. Substituting *λ* = *iω* in Eq. (45) and separating real and imaginary parts yields (*j* ∈ *I*_+_):

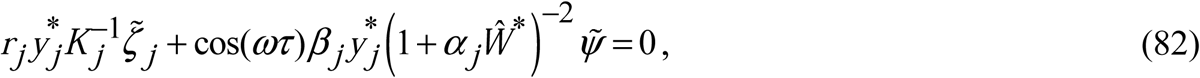

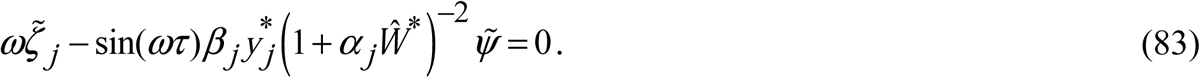

Since sin(*ωτ*) = 0, cos(*ωτ*) = 1, from Eqs. (75), (76), (78), (79), (82), (83) it follows that nontrivial solution can exist only if *ω* = 0, i.e. the neutral stability state is not reachable and characteristic Eq. (61) does not have complex root *λ* with non-negative real part. Thus, if *I*_0_ ≠ ∅ and 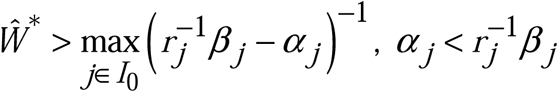, or if *I* =∅ the nontrivial equilibrium is unconditionally (i.e. for all *τ* > 0 [38]) locally asymptotically stable.

We can conclude that the digestion period of generalist consumer (time delay *τ*) does not cause local asymptotical instabilities of consumer population in the vicinity of nontrivial equilibria as one could expect. The results considered above are summarized in the following theorem.

#### Theorem 4.

Let coefficients of the system (1) – (6) satisfy conditions (7), (8), the nontrivial equilibrium 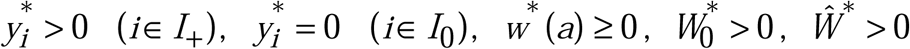 of autonomous system (1) - (5) is a solution of equation *R*(*C*(*y*\*(*Ŵ*\*),*Ŵ*\*)) = 1 with restrictions 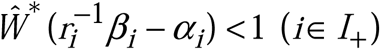 satisfied Eqs. (11), (12), (20), (21).

This equilibrium is unstable for all *τ* > 0 if *I*_0_ ≠∅ and

i. Ǝ*i*∈ *I*_0_ for which 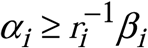, or
ii. 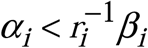 for all *i*∈ *I*_0_, and 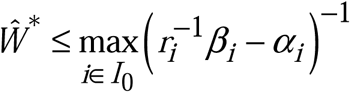. This equilibrium is unconditionally locally asymptotically stable (for all *τ* > 0) if
iii. 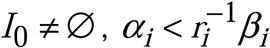 for all *i*∈ *I*_0_ and 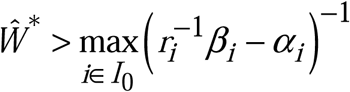, or
iv. *I*_0_ =∅.

Digestion period of generalist consumer *τ* does not cause local asymptotical instabilities of consumer population at the nontrivial equilibria.

#### Remark 1.

According to the statement (iv) of Theorem 4 the nontrivial equilibrium with non-depleted food patches (*I*_0_ =∅) is always locally asymptotically stable. That means that there exists the balance between resource growing, demographical process of consumer population and their consumption regime (within framework of the considered model) which guarantees the steady coexistence of all non-depleted resource patches with non-trivial consumer population. The local asymptotic stability of nontrivial equilibria of nonlinear age-structured models with density dependent fertility and death rates is well predicted by the partial derivative of basic reproduction number ([8], [13]). Such stability indicator of non-trivial equilibrium with non-depleted patches of system (1) – (6) has the form:

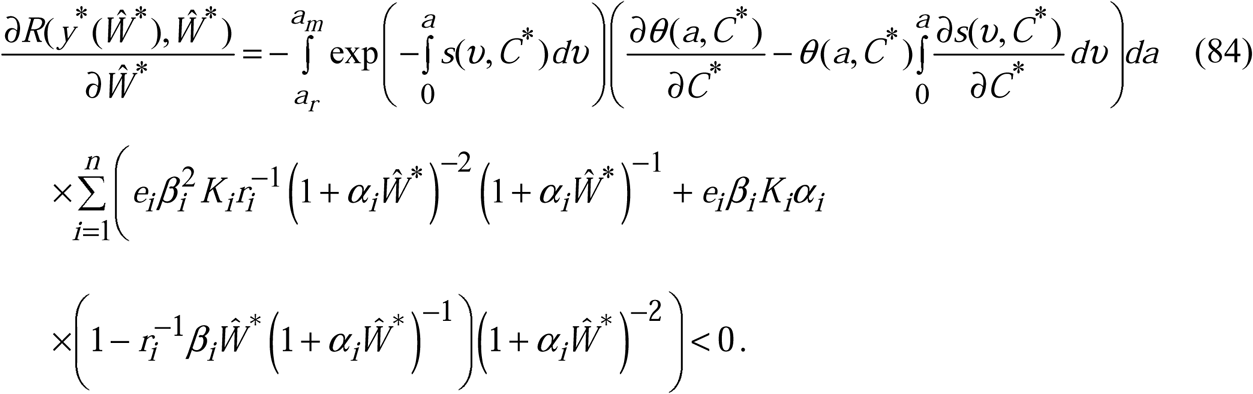

The negative value of this expression indicates the local asymptotic stability of the non-trivial equilibrium (see [8], [13]) with non-depleted patches that confirms the statement (iv) of Theorem 4. Since we could not adapt this indicator for analysis of local asymptotic stability of nontrivial equilibria with depleted patches (cases (i), (ii), (iii) of Theorem 4), anywhere further we will use the conditions of local asymptotic stability given in Theorem 4.

## 5. Numerical experiments

### 5.1. Parameterization of autonomous system (1) – (5)

We assume that the consumer fertility rate is an increasing with saturation monotone function and death rate is a decreasing with extinction monotone function of calorie intake rate satisfied Eqs. (7), (8). They are defined on the parametrized classes of algebraic functions:

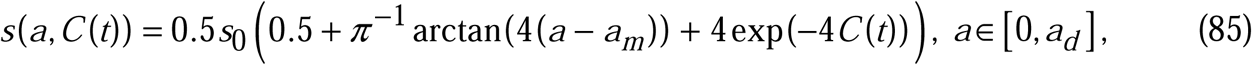

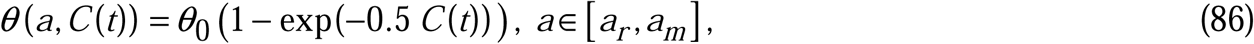

where *s*_0_, *θ* _0_ are given constants. The generalist consumer population is partitioned into three age-structured groups with young, matured and senile individuals with a number of individuals in each group 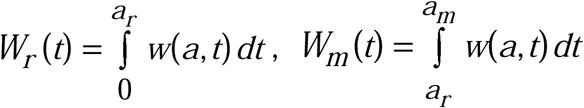 and 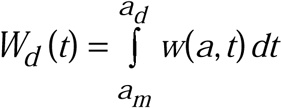, respectively. We use the following piece-constant function of resource intake weighted coefficient among age-structured groups (*γ*(*a*) ∈ *L*_2_ ([0, *a*_*d*_])):

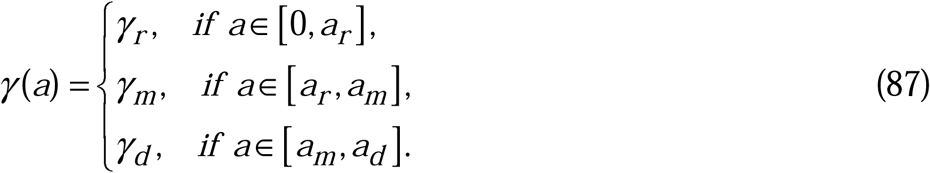

where 0 < *γ* _*r*_ < *γ* _*d*_ < *γ* _*m*_ < 1 - the set of constant dimensionless weights of resource intake for young, senile and matured consumers, respectively. The biggest value of *γ* _*m*_ in comparison with *γ* _*d*_ and *γ* _*r*_ means that one matured consumer takes more resource biomass than young or senile consumer.

For illustration of theoretical results obtained in Theorems 3 and 4, we consider the minimal set of three food resource patches in all experiments in the vicinities of stationary equilibria:

1. trivial and semi-trivial equilibria with small *τ* = 0,001*a*_*d*_ (Theorem 3);
2. trivial and semi-trivial equilibria with time delay from interval *τ* = 0,001*a*_*d*_, …, *τ* = 0,03*a*_*d*_, (Theorem 3);
3. positive equilibrium with one non-depleted patch, less active and more active depleted patches (statement (i) of Theorem 4);
4. positive equilibrium with one non-depleted patch and two less active depleted patches with 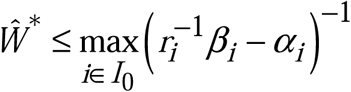 (statement (ii) of Theorem 4);
5. positive equilibrium with one non-depleted patch, two less active depleted patches with 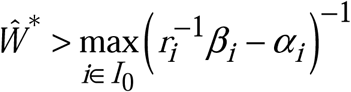 (statement (iii) of Theorem 4);
6. positive equilibrium with three non-depleted patches and nontrivial consumer population (statement (iv) of Theorem 4).

The values of coefficients of Eqs. (1), (6) and initial values (3), (4) vary in each experiment depending on the conditions of Theorems 3, 4.

### 5.2. The trivial and semi-trivial equilibria

The numerical algorithm based on the method of characteristics ([4] - [6]) is used here for study the dynamical regimes of autonomous system (1) – (5) in the vicinities of all equilibria considered in section 4.

In the first group of experiments we study the asymptotic behaviour of solutions in the vicinity of the trivial and semi-trivial equilibria with fixed small value of time delay *τ* = 0,001*a*_*d*_. The dynamics of mean resource density 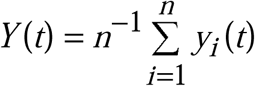, the quantity of consumers 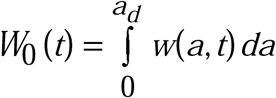 and the basic reproduction number *R*(*t*) are shown in Figs. 1a – 1c. The numerical simulations illustrate the statements of Theorem 3:

**Fig.1.**
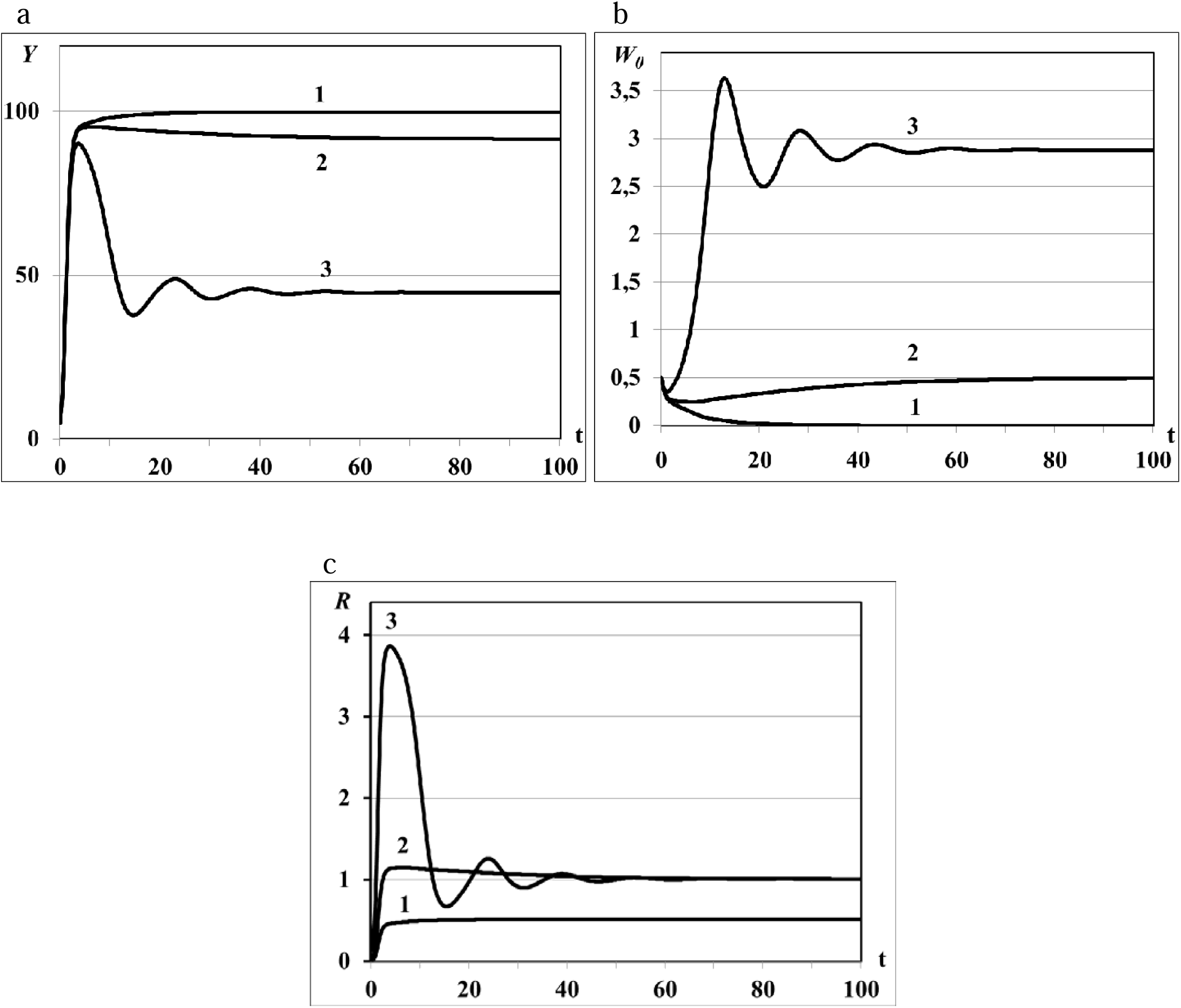
Graphs of asymptotic convergence of solutions with *R*(*K*) < 1 (curve 1), *R*(*K*) > 1 (curve 2), *R*(*K*) >> 1 (curve 3).

i. the trivial equilibrium is unstable (all curves in Figs.1.a – 1.c), *Y*(*t*) and *W*_0_(*t*) evolve to the semi-trivial equilibrium (curves 1 in Figs.1.a – 1.b, consumer population becomes extinct while the resource biomass in all patches saturates, *R*(*K*) < 1) or evolve to the nontrivial equilibrium (curves 2 and 3 in Figs.1.a – 1.b, *R*(*K*) > 1);
ii. the semi-trivial equilibrium is asymptotically stable with *R*(*K*) < 1 (curves 1 in Fig.1.a – 1.c) whereas it is unstable with *R*(*K*) > 1 (graphs 2 in Figs.1.a – 1.c).

For the very large basic reproduction number *R*(*K*) >> 1 obtained with large value of *θ*_0_(Eq.(86)) we observe the oscillatory regime of the system with asymptotic convergence of solution to the steady state (curve 3 in Figs.1a – 1c). The existence of such periodic solutions of some Lotka-Volterra prey-predator models was proved in theoretical work [45] and was observed in numerical experiments in [6], [7].

Further increasing of basic reproduction number by parameter *θ*_0_ causes the consumer population outbreaks (special dynamical regimes of population, see [1], [7], [9]). The pulse sequence or sequence of outbreaks (Figs. 2a – 2c) of consumers population and resource densities describe the quasi-periodic dynamical regime in the vicinities of the trivial and semi-trivial equilibria. The fast growing of consumer population is accompanied by the huge resource consumption, and as a consequence, by resource extinction and following decreasing of consumer population density to the minimal but not critical values. System moves to the trivial equilibrium from the vicinity of unstable semi-trivial equilibrium (*R*(*K*) >> 1, statement (ii) of Theorem 2). But, since the trivial equilibrium is unstable too (statement (i) of Theorem 2), and the minimal number of consumers is sufficient for the following renewal of population, system moves to the semi-trivial equilibrium again. This process is repeated at quasi-periodic time intervals and results in the pulse sequence of consumer population and resource densities. The same regimes were obtained in work [6] for the nonlinear age-structured model of population dynamics with density-dependent delayed death rate only for the big values of delay parameter and/or for the periodic time-dependent death and fertility rates. Since in this experiment the impact of the time delay parameter is insignificant and all coefficients of model are time-independent the dynamical regimes of periodic outbreaks are result of the repeating dynamical interaction between total resource consumption and its renewing from the one hand and consumer population growth and extinction from the other hand.

**Fig.2.**
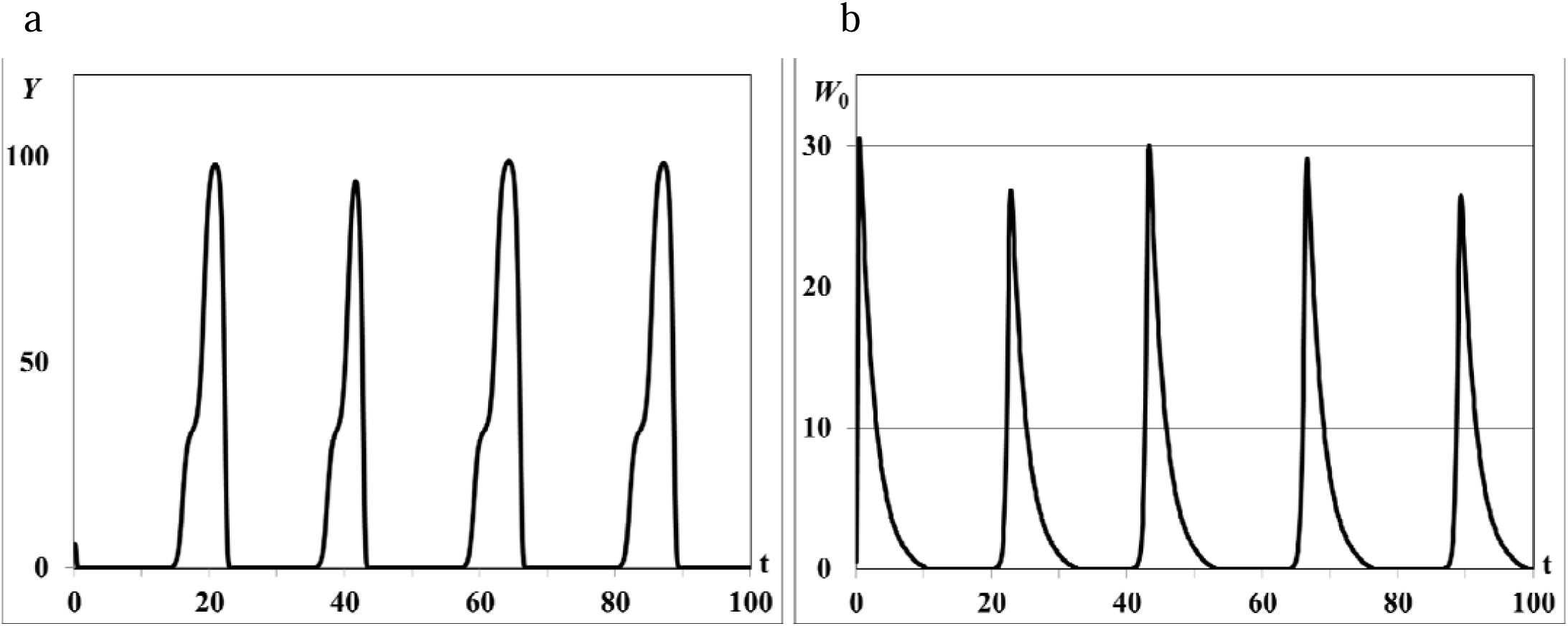
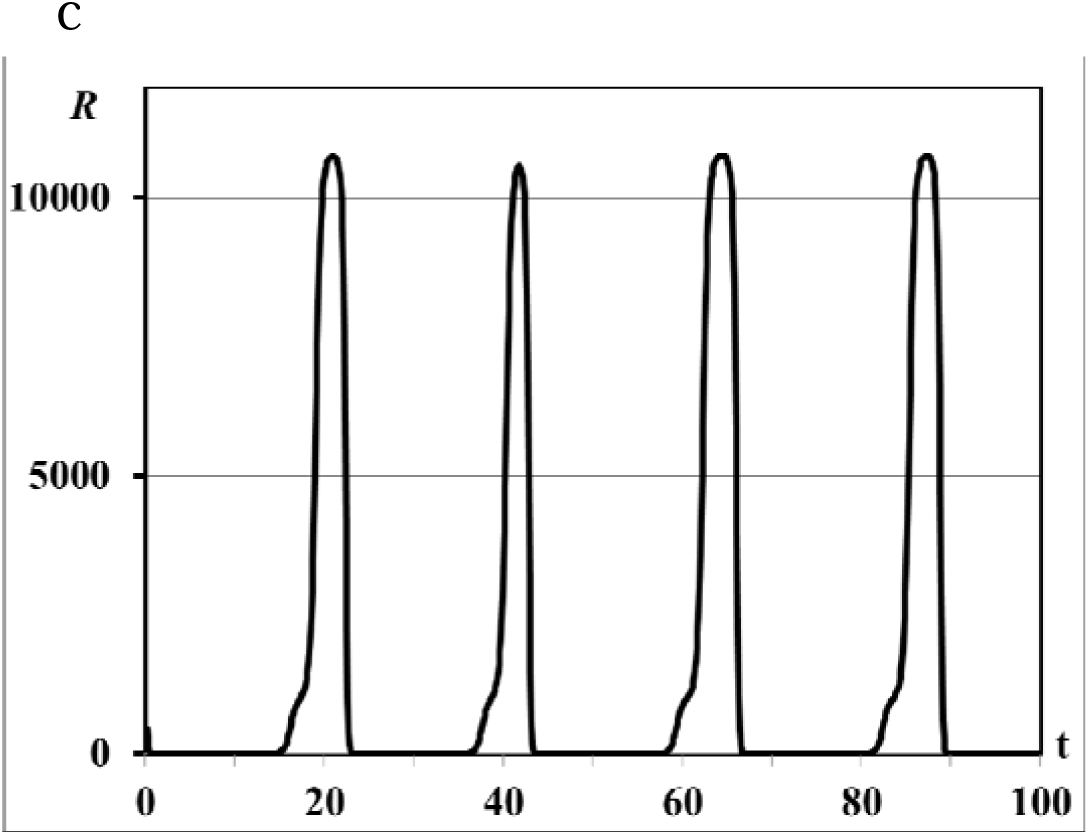
Graphs of periodic population outbreaks with *R*(*K*) >> 1, *τ* = 0,001*a*_*d*_.

Further increasing of parameter *θ*_0_ leads to the consumer population outbreaks of the single pulse form (Figs. 3a, 3b). The same rapid consumer population growth like in the previous experiment is accompanied by the huge resource consumption and resource extinction but with following decreasing of consumer population density up to the critical values when population is not able to renew the reproduction and becomes fully extinct (see Remark 1 to Theorem 3). The food resources in all patches saturate with time and system evolves eventually to the asymptotically stable semi-trivial equilibrium (Figs. 3a).

**Fig.3.**
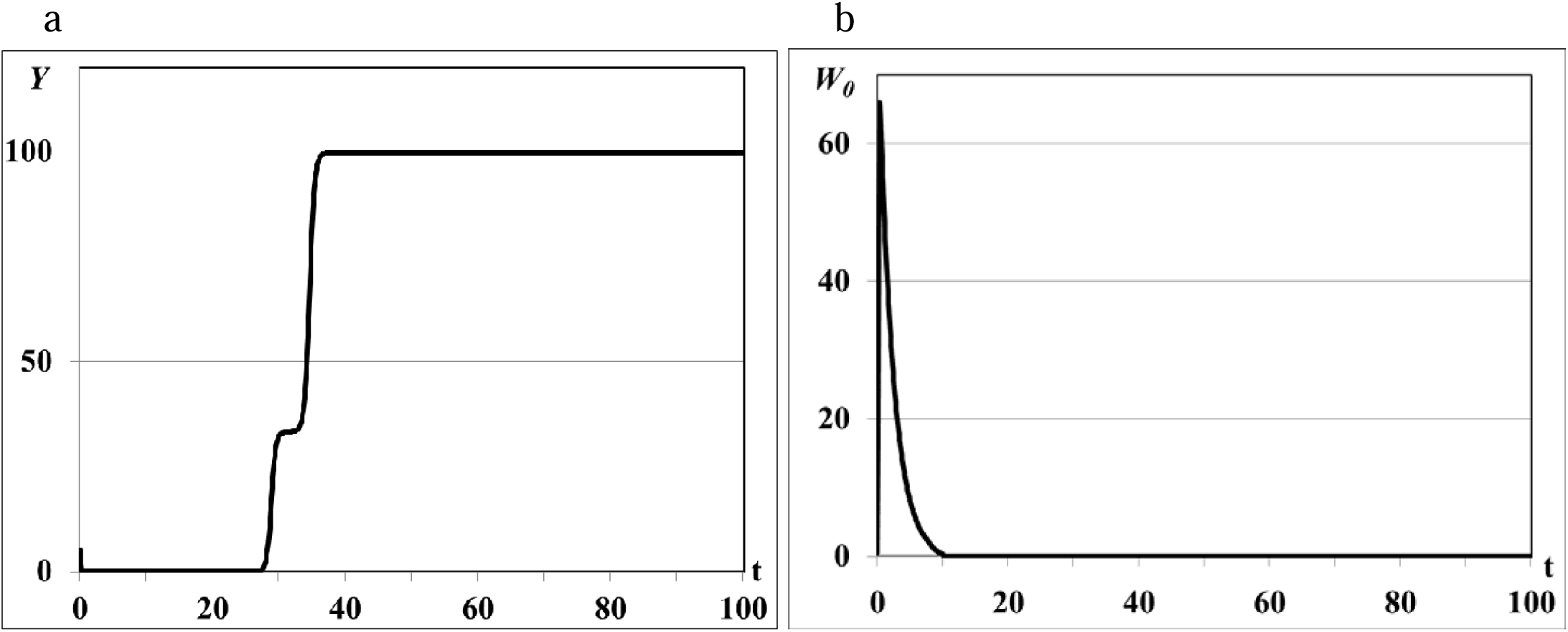
Graphs of population single outbreak and extinction, *τ* = 0,001*a*_*d*_.

In the second group of experiments we study the dynamical regimes of autonomous system (1) - (5) in the vicinity of the trivial and semi-trivial equilibria with different values of time delay from interval *τ* = 0,001*a*_*d*_, …, *τ* = 0,03*a*_*d*_, and large basic reproduction number *R*(*K*) >> 1. Solution of autonomous system (1) – (5) remains stable oscillated with bounded magnitude in the vicinity of the semi-trivial equilibrium in all experiments with different value of time delay (Figs.4 – 6). For the small value of *τ* = 0,001*a*_*d*_ the trajectories of system have the magnitude with exctincted oscillations and converge to the positive equilibrium (curve 3 in Figs.1a-1c). The bigger value of *τ* = 0,005*a*_*d*_ causes the periodic dynamics of *Y*(*t*) and *W*_0_(*t*) with bigger magnitudes, shown in Figs. 4a, 4b. Further increasing of time delay leads to the periodic outbreakes of *Y*(*t*) and *W*_0_(*t*) with *τ* = 0,015*a*_*d*_ (Figs.5a, 5b) and single outbreak of *W*_0_(*t*) with saturating *Y*(*t*) with *τ* = 0,03*a*_*d*_ (Figs.6a, 6b). The graphs shown in Figs.4 and 5 correspond to the regimes of population outbreaks obtained in the previous experiments (Figs. 2, 3). Similar dynamical regimes were obtained and described in works [6], [7] for the age-structured model with density-dependent delayed death rate and discussed in work [8] for the two-compartment age-structured model of locust population dynamics.

**Fig.4.**
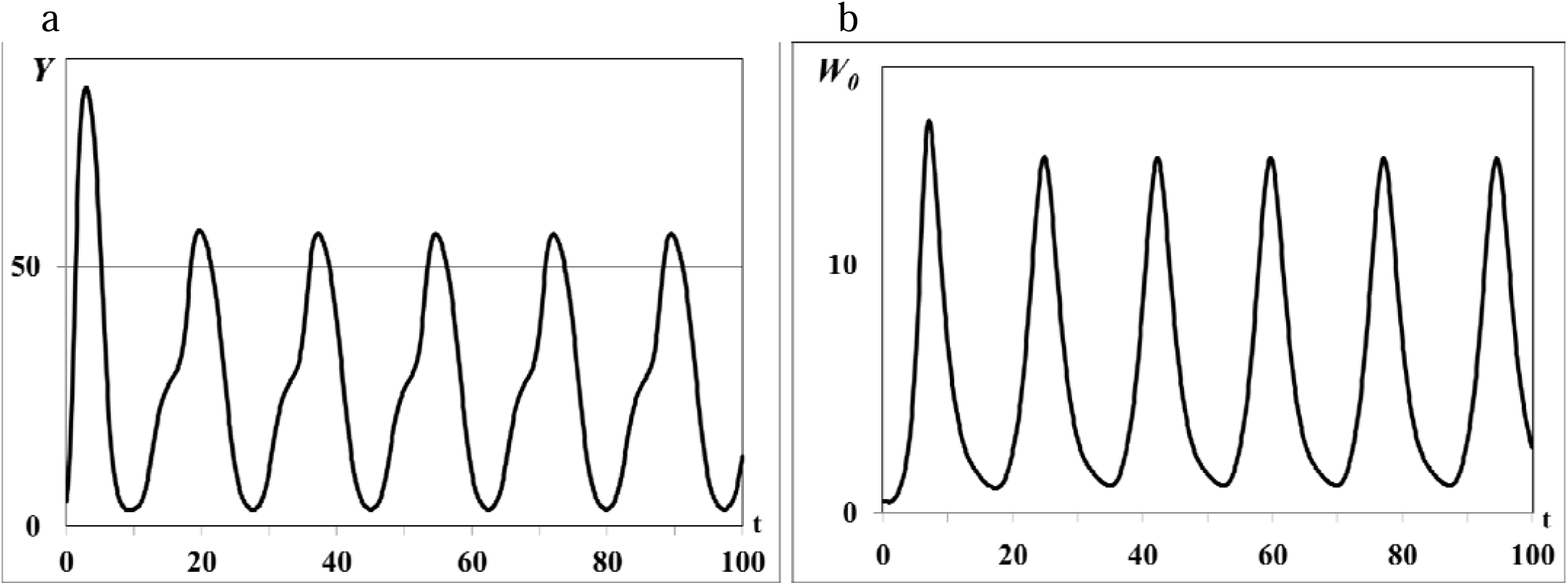
Graphs of periodic dynamics of *Y*(*t*) and *W*_0_(*t*), *τ* = 0,005*a*_*d*_.

**Fig.5.**
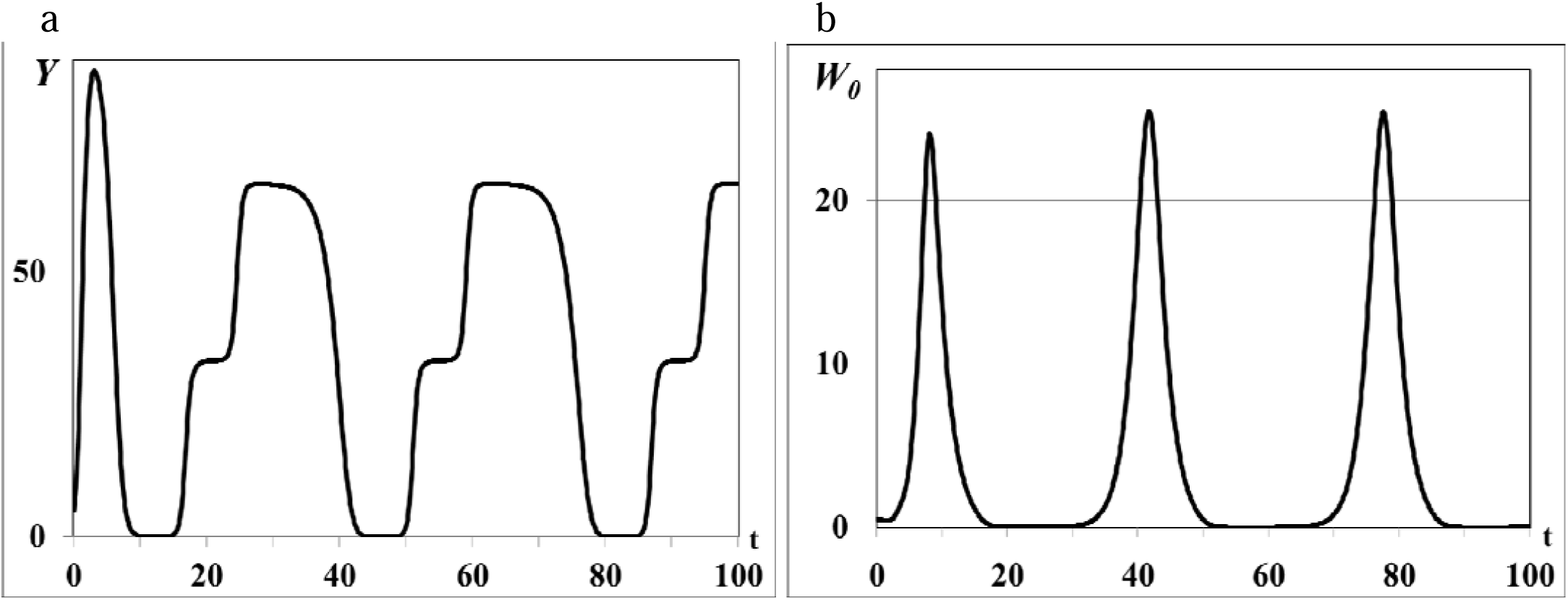
Graphs of periodic outbreaks of *Y*(*t*) and *W*_0_(*t*), *τ* = 0,015*a*_*d*_.

**Fig.6.**
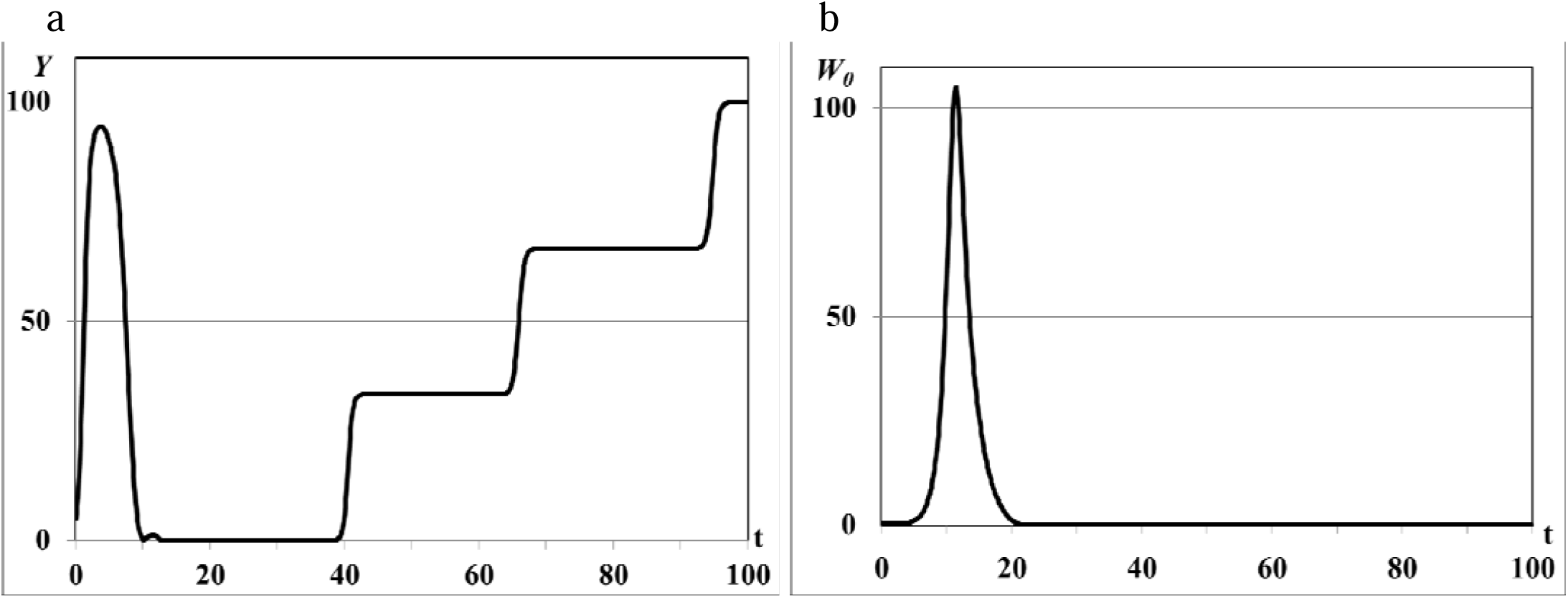
Graphs of asymptotic saturation of *Y*(*t*) (a), of single outbreak of *W*_0_(*t*) *τ* = 0,03*a*_*d*_.

The results of this group of experiments illustrate also that digestion period of generalist consumer *τ* does not lead to the high-frequency oscillations of solution in the vicinity of the semi-trivial equilibrium known as deterministic chaos.

### 5.3. The nontrivial equilibria

The third group of numerical experiments focuses on the study of asymptotic behaviour of solutions in the vicinity of the non-trivial equilibrium where some patches are depleted, i.e. 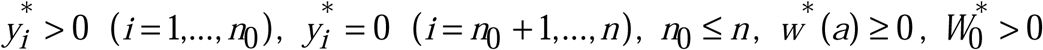. In all experiments *n* = 3, but *n*_0_ varies in each experiment depending from the condition of Theorem 4. For the fixed values of *n, n*_0_ the root of transcendental equation *R*(*C*(*y*\*(*Ŵ*\*),*Ŵ*\*)) = 1 is defined numerically by bisection method with restrictions 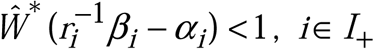. The equilibrium values 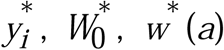, are defined by Eqs. (11), (20), (21) respectively. The value of *Ŵ*\* is used also in figures for illustration of asymptotic convergence of trajectories to the equilibrium.

In first experiment 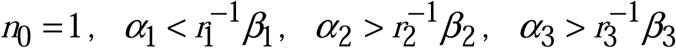. In equation *R*(*C*(*y*\*(*Ŵ*\*),*Ŵ*\*)) = 1 we use 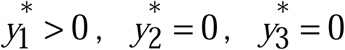, for which 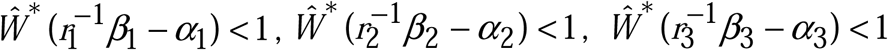 (case (i) of Theorem 4). The dynamics of resource densities *y*_*i*_(*t*) (*i* = 1, 2,3) and consumer weighted quantity *Ŵ*(*t*) are shown in Figs. 7a, 7b. The density of first food patch *y*_1_(*t*) does not evolve to the equilibrium 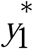 (curve 1 in Fig. 7a), patches *y*_2_(*t*) and *y*_3_(*t*) are not depleted (curves 2 and 3 in Fig. 7a) and consumer weighted quantity *Ŵ*(*t*) does not evolve to the equilibrium *Ŵ*\*(Fig. 7b), i.e. the nontrivial equilibrium is unstable.

**Fig.7.**
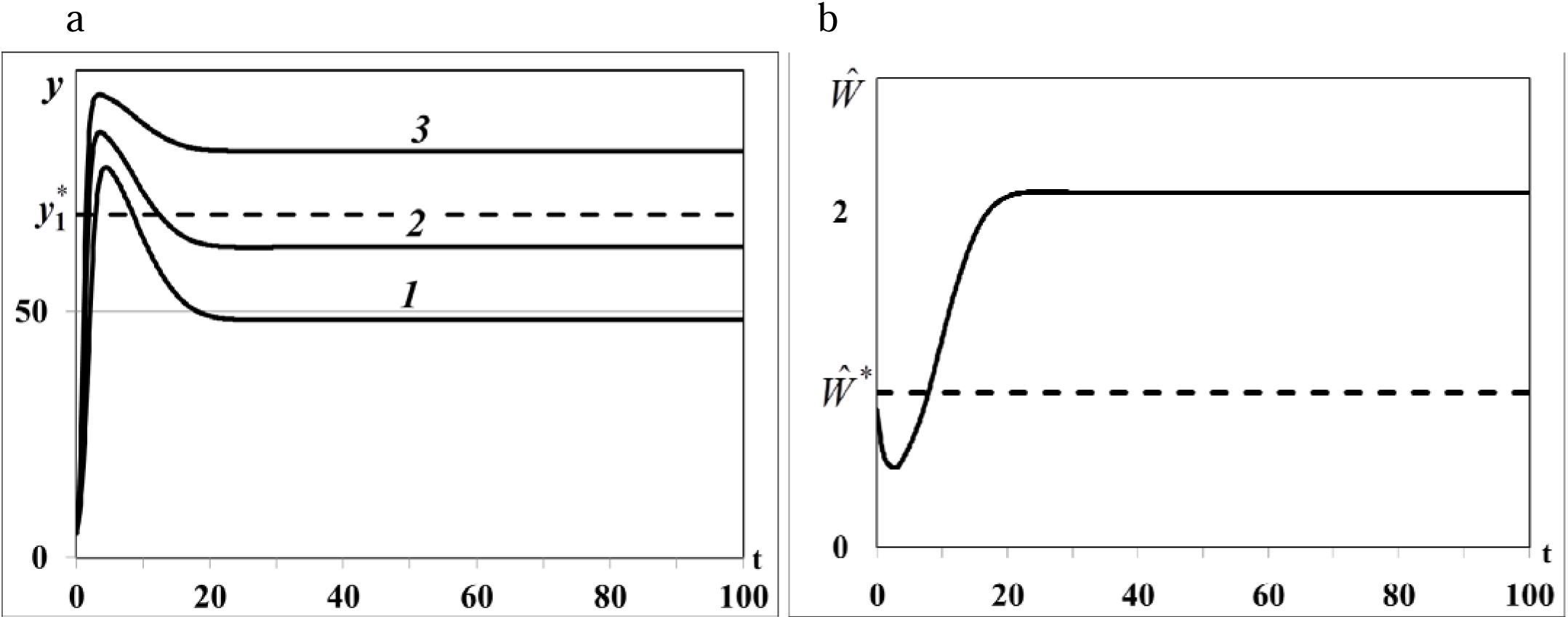
a. Case (i). Resource densities *y*_*i*_(*t*) (*i* = 1, 2,3). b. Weighted number of consumers *Ŵ*(*t*).

In second experiment 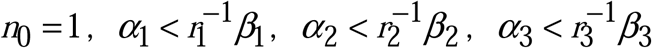. In equation *R*(*C*(*y*\*(*Ŵ*\*),*Ŵ*\*)) = 1 we use 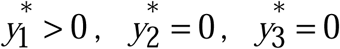, for which 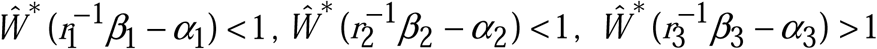 (case (ii) of Theorem 4). The density of first food patch *y*_1_(*t*) does not evolve to the equilibrium 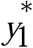 (curve 1 in Fig. 8a), second patch is not depleted (*y*_2_(*t*), curve 2 in Fig.8a), *y*_3_(*t*) converges to the trivial equilibrium and becomes depleted (curve 3 in Fig.8a) and consumer weighted quantity *Ŵ*(*t*) does not evolve to the equilibrium *Ŵ*\* (Fig. 8b), i.e. the nontrivial equilibrium of food web is unstable.

**Fig.8.**
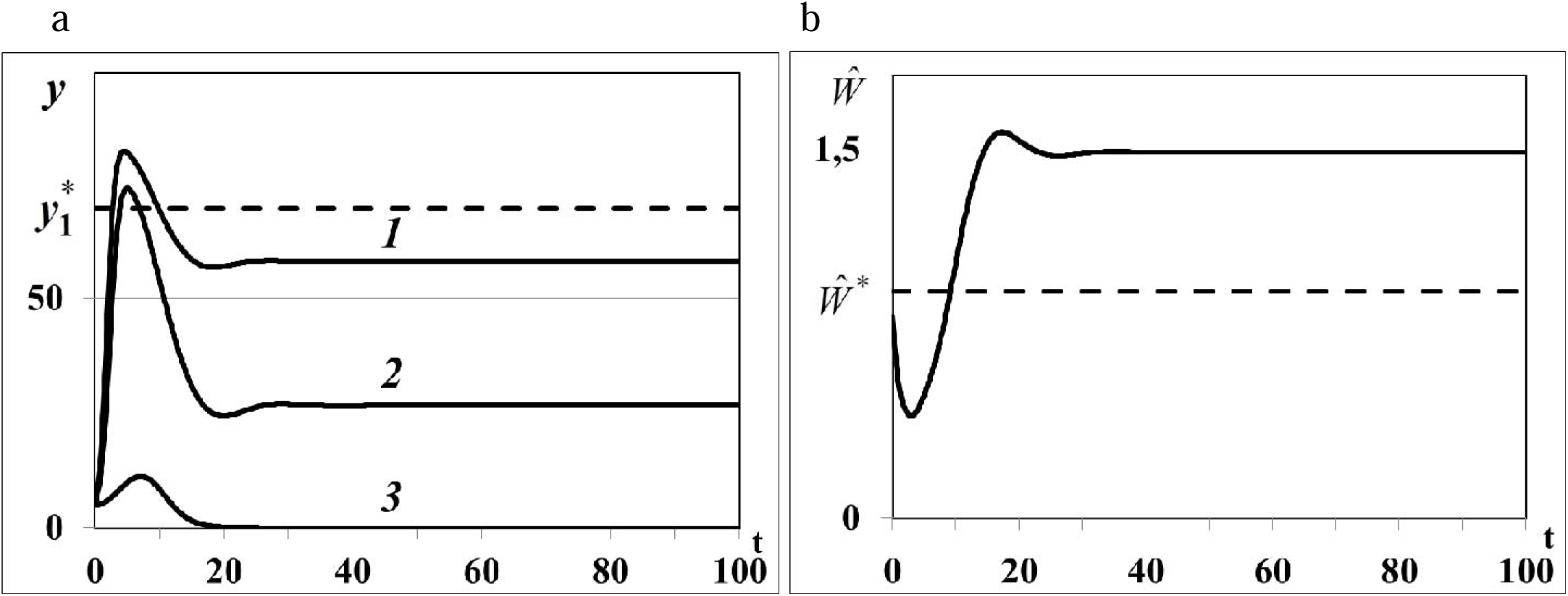
a. Case (ii). Resource densities *y*_*i*_(*t*) (*i* = 1, 2,3). b. Weighted number of consumers *Ŵ*(*t*).

In third experiment 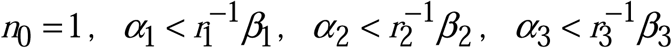. In equation *R*(*C*(*y*\*(*Ŵ*\*),*Ŵ*\*)) = 1 we use 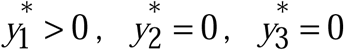, for which 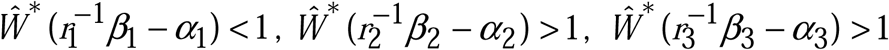 (case (iii) of Theorem 4). The density of first food patch *y*_1_(*t*) in this case evolves to the positive equilibrium 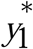 (curve 1 in Fig. 8a), the densities of the other patches *y*_2_(*t*) and *y*_3_(*t*) evolve to the trivial equilibrium 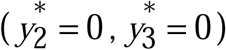, and become depleted (curves 2 and 3 in Fig. 8a). Consumer weighted quantity *Ŵ*(*t*) evolves to the positive equilibrium *Ŵ*\* (Fig. 8b), i.e. the nontrivial equilibrium with one non-depleted patch and two depleted patches is locally asymptotically stable.

In the last forth experiment 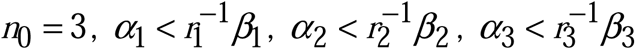. In equation *R*(*C*(*y*\*(*Ŵ*\*),*Ŵ*\*)) = 1 we use 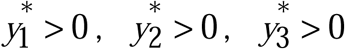, for which 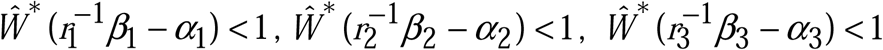 (case (iv) of Theorem 4). The densities of food patches *y*_*i*_(*t*) evolve to the corresponding positive equilibria 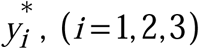 (curves 1, 2, 3 in Fig. 10a), consumer weighted quantity *Ŵ*(*t*) evolves to the positive equilibrium *Ŵ*\* (Fig. 10b), i.e. the nontrivial equilibrium with all non-depleted patches is locally asymptotically stable.

**Fig.9.**
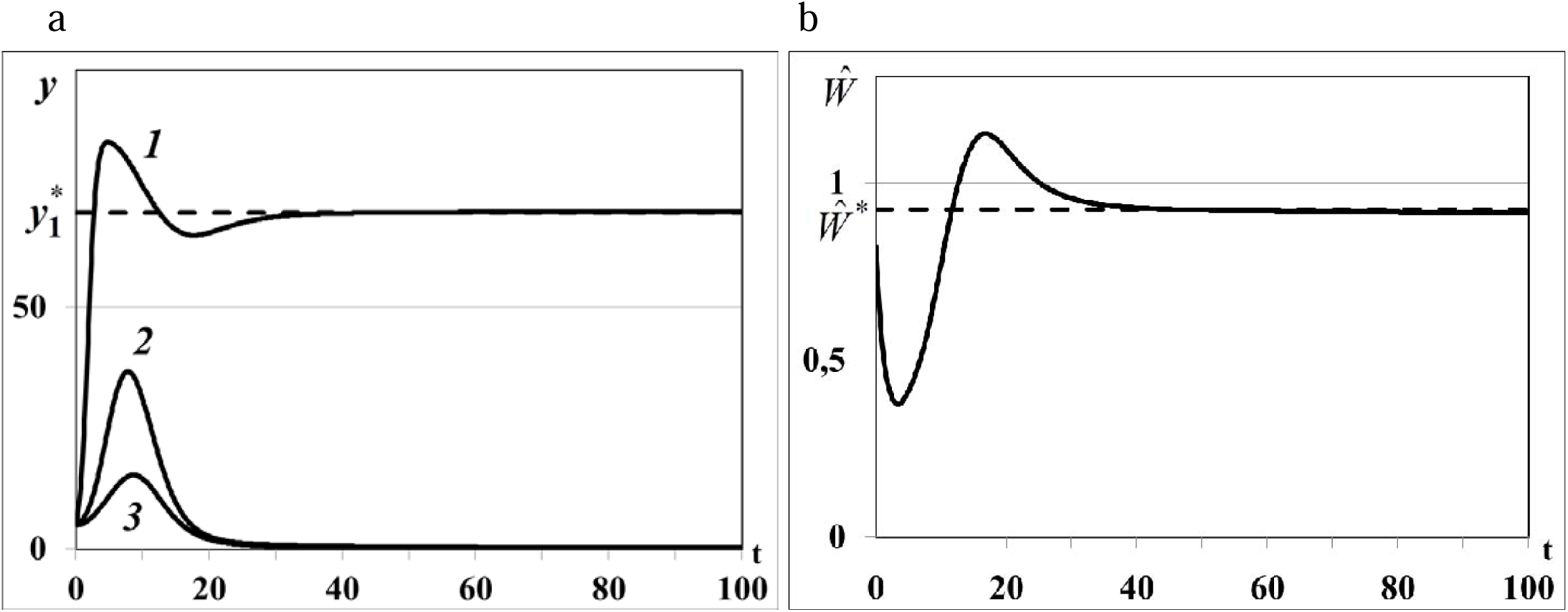
a. Case (iii). Resource densities *y*_*i*_(*t*) (*i* = 1, 2,3). b. Weighted number of consumers *Ŵ*(*t*).

**Fig.10.**
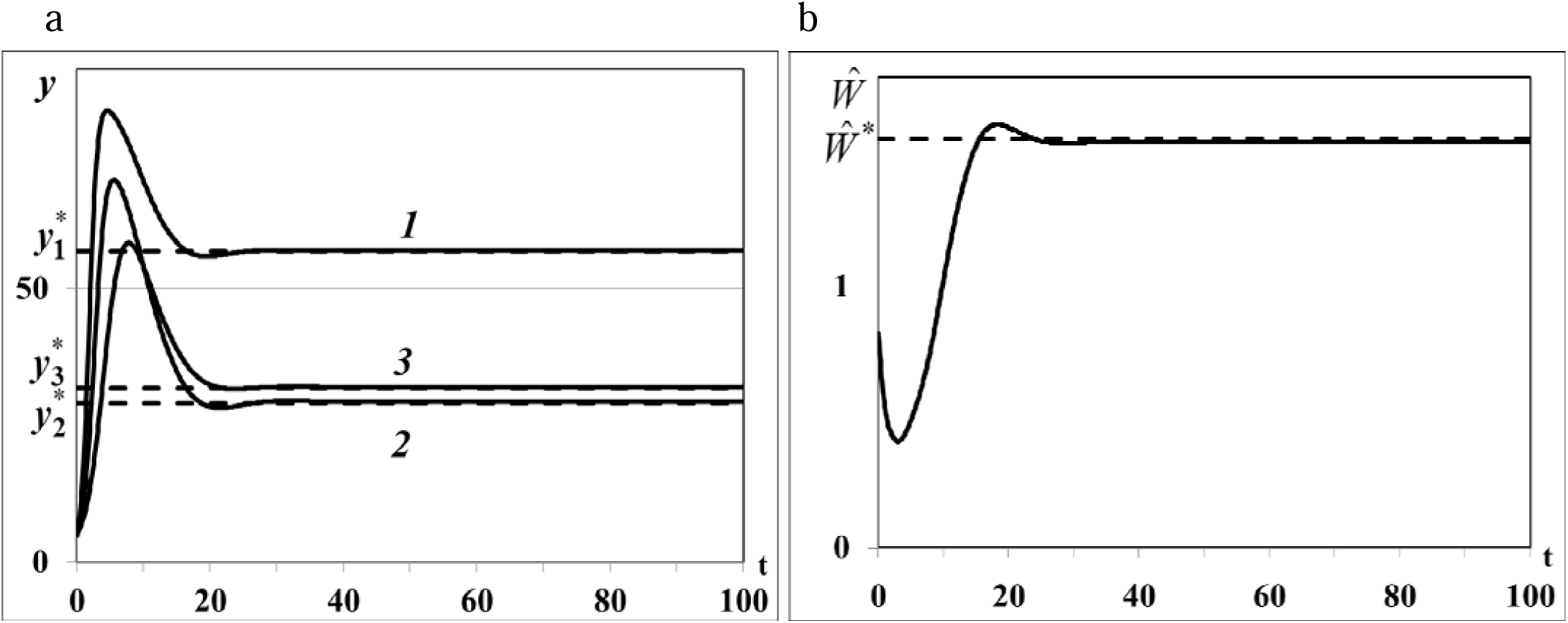
a. Case (iv). Resource densities *y*_*i*_(*t*) (*i* = 1, 2,3). b. Weighted number of consumers *Ŵ*(*t*).

The numerical results of third and fourth experiments illustrate the statements (iii) and (iv) of Theorem 4. In third experiment the nontrivial equilibrium with one non-depleted patch and two depleted patches (*I*_0_ ≠∅) and in forth experiment the nontrivial equilibrium with three non-depleted patches (*I*_0_ =∅) are locally asymptotically stable (Figs.9, 10). Overall, we can conclude, that all statements of Theorem 3 and Theorem 4 correctly predict the asymptotic stability or instability of trivial, semi-trivial and non-trivial equilibria of system (1) – (5) in all numerical experiments.

## 6 Conclusions and discussions

In this work it was studied an autonomous system – a resource-consumer model in a heterogenous environment consisting of several food patches with active resource. Food resources do not disperse between patches, while consumers do disperse. The model of food resources is unstructured while the model of consumer population is age-structured. The relationship between the consumed food resource and consumer demographic parameters (fertility and death rates) is modelled by means of a calorie intake rate that describes the amount of energy obtained by consumer at a given age from all food patches per unit of time. In biological applications calorie intake rate can be obtained from the observations, or foraging experiments focusing on age-structured consumer behaviour. The consumer calorie intake rate is proportional to the saturated intake rate (where the coefficient of saturation is a behavioural parameter of food resource) and depends from the time period a consumer needs to handle and digest a unit of resource (delayed parameter). Thus, the model considered in this paper extends the classic apparent competition models [24] – [27], [30] to a structured consumer population with time delay and active food resources.

All types of possible equilibria: trivial, semi-trivial (with saturated food resource densities and extinct density of consumer population), and non-trivial equilibria with depleted and non-depleted patches and positive density of consumer population were studied. In all theorems, it was used a new condition of sign-preserving partial derivatives of calorie intake rate-dependent fertility and mortality rates of consumer: 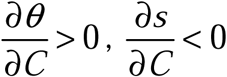. The trivial and semi-trivial equilibria of the nonlinear autonomous system always exist while the non-trivial equilibria exist if and only if the basic reproduction number of the consumer population *R* = 1 (Theorem 1). On the basis of this result it was obtained in Theorem 2 new and constructive sufficient conditions on coefficients of the autonomous system (on the basis of ideas and results from [14]) for existence of nontrivial equilibria with depleted and non-depleted foraging patches and positive density of consumer population.

The conditions of local asymptotic stability/instability of trivial and semi-trivial equilibrium obtained in Theorem 3 were formulated in terms of the consumer’s basic reproduction number. Stability analysis of non-trivial equilibria was carried out on the basis of perturbation theory and linearization method. Unfortunately, the well-known stability indicator of equilibria of nonlinear age-structured models – partial derivative of density-dependent basic reproduction number of consumer population ([8], [13]) cannot be used for stability analysis of nontrivial equilibria of resource-consumer model with depleted patches. Instead of it the explicit conditions on coefficients of system were obtained in Theorem 4 for instability/local asymptotic stability of nontrivial equilibria with several or without depleted resource patches. In all theorems it was rigorously proved that the time-delay parameter - the consumer’s digestion period does not cause local asymptotical instabilities of consumer population at the trivial, semi-trivial and nontrivial equilibria.

The dynamical regimes of autonomous system with the different values of time-delay parameter were studied in numerical experiments for illustration of results obtained in theorems. In the 1-st and 2-nd groups of experiments the local asymptotic stability/instability of the trivial and semi-trivial equilibria, consumer population outbreaks, extinct and nonextinct quasi-periodic dynamic regimes were obtained for the different values of the time delay parameter. The processes of resource handle and food digestion are inherent for all biological organisms, although the value of handling and digestion period can significantly differ among species. Numerical experiments showed that digestion period of generalist consumer *τ* does not cause the local asymptotical instabilities or high-frequency oscillations (deterministic chaos) of consumer population in the vicinity of semi-trivial equilibrium. All numerical experiments were carried out in correspondence with the conditions formulated in Theorems 1 - 3.

In the 3-rd and 4-th groups of experiments we study the local asymptotic stability/instability of the nontrivial equilibria of system with one non-depleted and two depleted resource patches (3-rd group) and three non-depleted resource patches (4-th group) and one generalist consumer that illustrated four statements of Theorem 4. The most important in practice theoretical conclusion of Theorem 4 confirmed by numerical results is that if there exists the non-trivial equilibrium - positive solution of equation *R*(*C*(*y*\*(*Ŵ*\*),*Ŵ*\*)) = 1, which satisfies condition 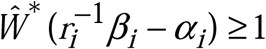 for all depleted patches (*i*∈ *I*_0_) and 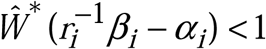 for all non-depleted patches (*i*∈ *I*_+_) such equilibrium is always locally asymptotically stable. The coefficient of saturation (behavioural characteristic of a food resource) *α*_*i*_ plays an important role in this criterion: if 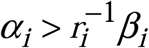 the corresponding *i* -th patch cannot be depleted in the asymptotically stable equilibrium of food web. Thus, the numerical results illustrated and confirmed all statements of Theorems 1 – 4 obtained in this paper.

## ACKNOWLEDGEMENTS

I would like to acknowledge professor Vlastimil Krivan for the helpful and valuable comments and suggestions to the topic of manuscript.

